# A General Analytic Approach to Predicting the Best Antibiotic Dosing Regimen

**DOI:** 10.1101/2025.09.13.676026

**Authors:** Leah Childers, Pia Abel zur Wiesch, Jessica M. Conway

**Affiliations:** Department of Mathematics, Pennsylvania State University, University Park PA, U.S.A; Department of Pharmacy, UiT – The Arctic University of Norway, Tromsø, Norway

**Keywords:** antibiotic dosing, pharmacokinetics and pharmacodynamics (PK/PD), dose-response curves, mathematical modeling, antimicrobial resistance (AMR)

## Abstract

Determining optimal antibiotic dosing strategies is complex. Clinically, some antibiotics work best in continuous low doses, while others require high repeated pulses. However, a rational understanding of the best approach depending on the specific pairing of antibiotic and bacterial species remains unclear. Using mathematical models, we analyze bacterial populations under two strategies – constant concentration and repeated dosing – given fixed pharmacodynamic and pharmacokinetic properties. Our results reveal that the shape of the dose-response curve, which measures bacterial net growth rate against antibiotic concentration, is crucial. Specifically, its concavity determines the best strategy. In cases where the curve exhibits multiple concavities, additional factors such as tolerable dosing range influence the regimen. These findings challenge the universal application of “hit hard and hit early,” as some recommended schedules include lower, constant doses. This work contributes to the literature on rational antibiotic prescription, aiming to minimize antibiotic use and combat antimicrobial resistance.

## 1 Introduction

Since the discovery of penicillin in 1928, antibiotics have been effective in eliminating numerous bacterial infections. However, irrational antibiotic usage threatens to undermine this success at both the individual and population levels. As expanded upon below, underdosing can lead to treatment failure while overdosing can cause adverse side effects; both under- and over-dosing can contribute to the development of antibiotic resistance [6, 19, 34]. Thus we want to expose a patient to as little antibiotic as possible, but as effectively as possible. In this paper, we fix a quantity of antibiotics and determine best candidate regimens that use the fixed quantity of antibiotics to minimize the bacterial population.

Historically, treatment regimens have been influenced by arbitrary or external factors, such as drug cost, manufacturing difficulty, or the convenience of the dosing schedule, rather than based entirely on the intrinsic pharmacokinetic (PK) and pharmacodynamic (PD) properties of the drug. For example, when treating tuberculosis with rifampin, the drug is often dosed at 600 mg daily due to historical expense concerns [37], although more recent studies suggest that higher doses may be more effective [27]. Van Ingen et al. found no clinical justification for the 600 mg dose when reviewing over 1,000 publications [37]. As drug price and availability become less restrictive, using the PK and PD properties to optimize dosing is crucial for minimizing short and long-term negative effects of antibiotics at both the individual and population levels, as well as minimizing cost of treatment [18], all while ensuring the treatment remains effective. At the individual level, balancing the negative effects of both high and low doses of antibiotics can alter the treatment outcome; allergic reactions [3], increased frequency and severity of side effects, and antibiotic resistance at the individual level can occur when antibiotics are prescribed frequently or in high doses[21]. At the population level, the irrational usage of antibiotics is a significant cause of the development of antibiotic resistance [6, 19], since exposure of bacteria to sublethal doses of antibiotics may increase the likelihood of emergent mutations leading to resistance [34]. Resistant infections are becoming more difficult, sometimes impossible, to treat, posing a global health crisis.

Currently, much is still unknown about what the best dosing strategy is for a given antibiotic and bacterial infection [18] in general. Progress made on this question has largely been case-by-case [16, 25, 26]. A recent 2023 review describing contemporary international antibiotic dosing practices suggests that while many changes have occurred since a similar survey (ADMIN-ICU [31]) in 2015, current practices still risk underdosing, overdosing, and the development of antimicrobial resistance [38]. This review also explains that the development of antibiotic resistance may increase when the drug’s PK/PD parameters are not considered. Corroborating the idea that much is still unknown about the theoretical best dosing strategy is that, in practice, outcomes have been mixed. Some research suggests well-timed repeated doses are more effective in some cases, such as against persister bacteria which can survive antibiotic treatment by going dormant [20, 28], while some suggest dosing at a constant concentration is more effective in some cases, such as in critically ill or immunocompromised patients [2, 10, 36].

Past studies have experimentally explored whether continuous (the same concentration for an extended period of time) or repeated (a certain amount being administered every *x* hours) dosing is more effective for particular antibiotic in very specific cases [10, 20, 36]. Many papers have acknowledged the ceiling effect of increasing antibiotic concentration [2, 14] and have classically modeled pharmacodynamics using Hill functions, noting that greater Hill coefficients result in steeper dose response curves [15, 25]. We will utilize Hill functions for most of the paper, so its properties, including shape, saturation threshold, and steepness, will play an impactful role in our analysis and predictions. Additionally, many specific PK/PD parameters and indices have been explored both experimentally and analytically [23], including the maximum effect of the drug (*E*_max_, PD), the concentration at which half the maximum effect is achieved (*EC*_50_, PD) [15], the minimum inhibitory concentration (MIC, PD) [25], time spent above MIC (%*T*_>MIC_, PK), the area under the drug concentration function (AUC, PK), the peak concentration of the drug (*C*_max_, PK), and ratios of these parameters [22]. While we used specific models and parameters to numerically demonstrate the results of this paper, our novel results are general and can be applied to any PK/PD model which satisfies our assumptions. Below we will describe our broad assumptions and setup, and a brief explanation of the nature of our results; after we will provide more context for past results from other modeling papers and how our results differ from them.

In the following we make the simplifying assumption that when we say “bacteria” in any context in this paper, we are assuming the bacteria population is of homogenous composition and that the antibiotic is equally effective against all of the bacteria in the population, and resistance is not a factor. We do this purposefully to develop the theoretical framework: Given a fixed antibiotic and its PK/PD properties for a given bacterial infection as well as a fixed total amount of antibiotic exposure in the body, the aim of this study is to identify the treatment strategy which minimizes the bacteria population. Fixing some notion of “total” is important when antibiotic or financial resources are limited. Specifically, we fix a repeated dosing regimen and compare its bacterial reduction to that of a regimen which holds the drug concentration constant during the entire treatment interval and has the same total antibiotic exposure (which is quantified as the area under the drug concentration function, or “AUC”) as the repeated dosing regimen.

Previous modeling papers have explored a similar model setup to ours, but they have asked different questions. For example, Katriel (2023) [17] aimed to identify the type of function describing antibiotic concentration which minimizes the AUC while achieving a specified reduction of the bacterial load. They prove that a continuous concentration dose given for a finite amount of time within the treatment length is the “ideal concentration profile” for minimizing the AUC. While Katriel focuses on minimizing the AUC, we focus on the most effective way to use a fixed AUC potentially representing a limited supply of antibiotics, essentially a dual question. Both Katriel and this present study contribute complementary advancements made to the rational prescription of antibiotics.

In the following, we aim to provide a framework for showing when constant concentration dosing is better, and when repeated dosing is better. To that end, we investigate two PK models of repeated dosing - one where we model the decay and accumulation of the drug (“Decay PK Model”), and one where we do not (“Step PK Model”), both coupled with a dose response curve and exponential bacterial growth rate model to test effectiveness. We prove a primary theorem for each of these PK models characterizing exactly at which concentrations the repeated dosing regimen performs better than the constant concentration regimen (and vice versa). We then illustrate our results using empirically found parameters for some common antibiotics. The results of our investigation challenge the conventional strategy of “hit hard and hit early,” suggesting that it may not always be the most effective approach. We show that sometimes, a low, constant concentration is more effective than a larger single dose. In those cases, one should “hit softly but hit constantly.”

## 2 Results

In this section we begin by describing our setup. In Sections 2.1 and 2.2 we prove analytical results about linear and nonlinear dose response curves respectively. While the results regarding linear dose response curves do not require any assumptions about the antibiotic concentration beyond comparing two regimens with the same AUC, we must be more specific about how we model pharmacokinetics when we investigate nonlinear dose response curves, which we do by introducing the “Step PK Model” in Section 2.2.2 and the “Decay PK Model” in Section 2.3. These models are described in detail these sections. In Section 3, we demonstrate our results and their implications numerically using empirically found parameters for some common antibiotics.

We conduct our analysis under the assumptions of a very simple unbounded bacterial growth model where the birth (replication) and death rates are combined into a single net growth rate [2, 15, 25]:

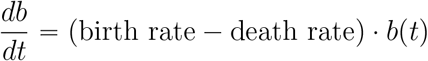

where *b* is the bacteria population (measured in any units, such as percentage of initial bacteria population). Let *R* = birth − death; then we can write this ODE as

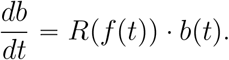

Note that antibiotics may modify the bacterial birth/replication rate, death rate, or both. The definitions of “bactericidal” and “bacteriostatic” antibiotics are often used to distinguish between these two effects, though these two terms do not have universally agreed upon definitions. In some modeling contexts, a descriptive definition is used: bacteriostatic antibiotics are those which inhibit growth without pushing the population into decline, while bactericidal antibiotics induce negative net growth rates [39]. However, in this paper we use mechanism-motivated definitions: bacteriostatic antibiotics inhibit bacterial growth by reducing the replication rate, while bactericidal antibiotics actively kill bacteria, increasing the death rate [24]. Both of these effects can lead to a negative net growth rate *R*.

The bacterial growth model we use here can be used to describe the net effects of both bacteriostatic and bactericidal antibiotics under these definitions, however the model does not separate these two effects. The antibiotic effect is then modeled the same for both types of antibiotics, simply as *R* = *R*(*f* (*t*)) where *f* (*t*) is the antibiotic concentration at time *t*. Thus *R* is the *dose-response curve*, and *R* can attain negative values for either case, which is consistent with the data generated by Regoes et al. to fit their pharmacodynamic curves [25] that we will use in Section 3. Example dose response curves are shown in Figures 1A and 2B. In this paper, we wish to compare the performance of different dosing regimens; to achieve this aim, we apply the following algorithm:

**Figure 1.**
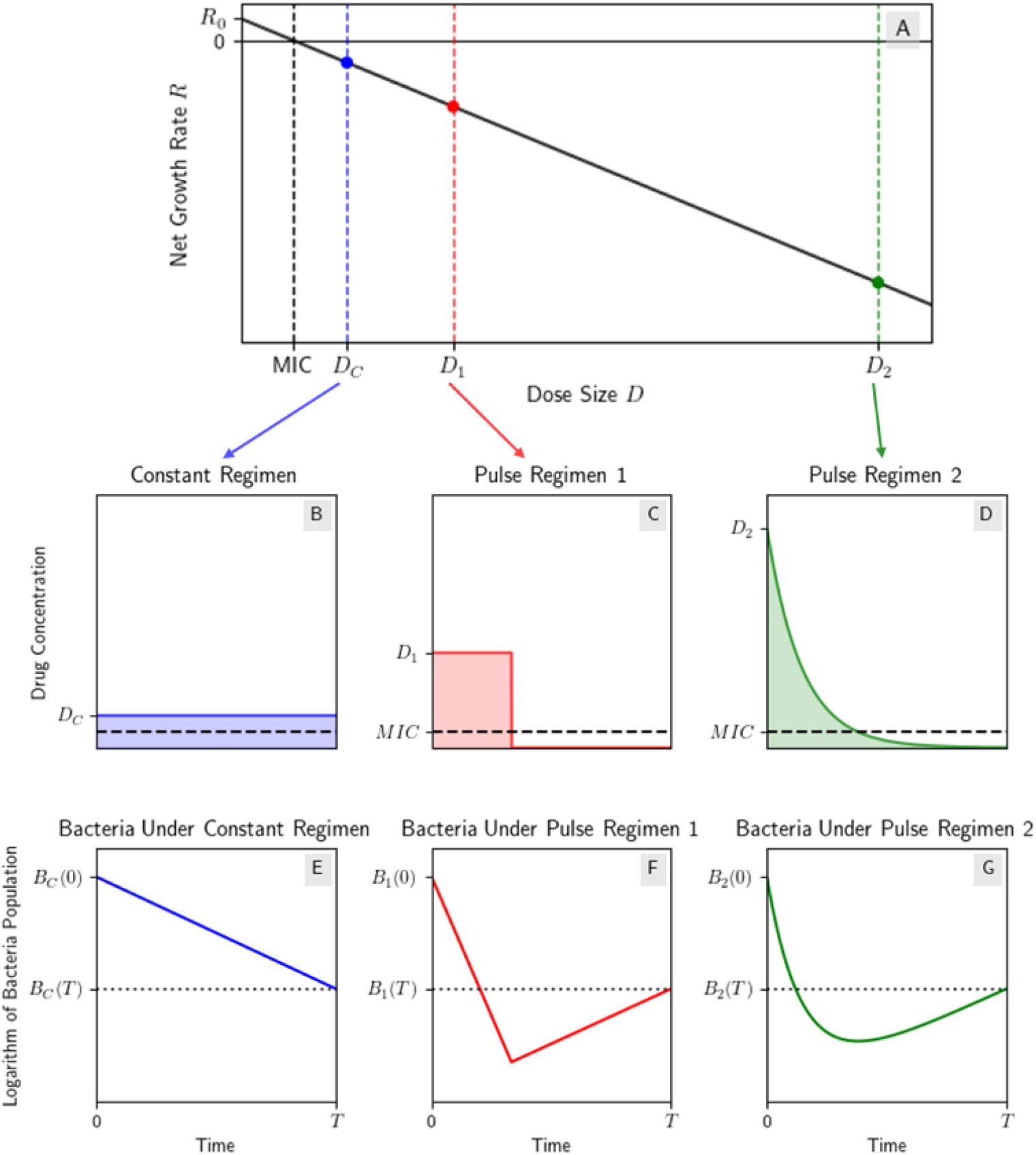
Dosing regimens with the same AUC under a linear dose response curve have equal end bacteria population. (A) Linear dose-response curve; (B) a CC regimen with drug concentration *D*_*C*_; (C) a “pulse” regimen using the step model from Section 2.2.2 with drug concentration *D*_1_ and the same AUC as the CC regimen; (D) a “pulse” regimen using the decay model from Section 2.3 with initial drug concentration *D*_2_ and the same AUC as the other two regimens; (E, F, G) the bacteria population *B*_*C*_, *B*_1_, *B*_2_ under the linear PD model using the three different dosing regimens respectively. Note that *B*_*C*_(*T*) = *B*_1_(*T*) = *B*_2_(*T*) as expected from Theorem 1. To reproduce result illustrated here, see Appendix F.

**Figure 2.**
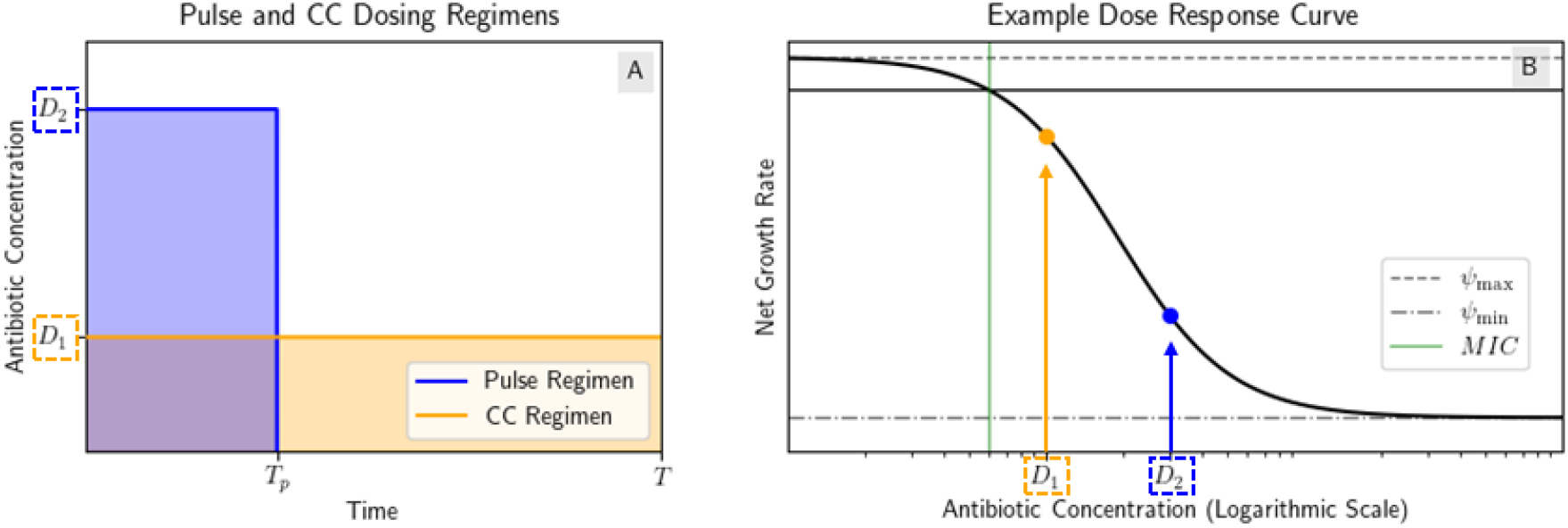
Example of step model regimens with the same AUC. (A) Pulse and CC regimens. The CC regimen has drug concentration *D*_1_ over the whole treatment interval [0, *T*] and the pulse regimen has drug concentration *D*_2_ over the interval [0, *T*_*p*_] and zero drug concentration over the interval [*T*_*p*_, *T*]. Both regimens have the same AUC. (B) Example dose response curve with the drug concentrations of the dosing regimens marked.

### Algorithm 1 Comparison of Dosing Regimens

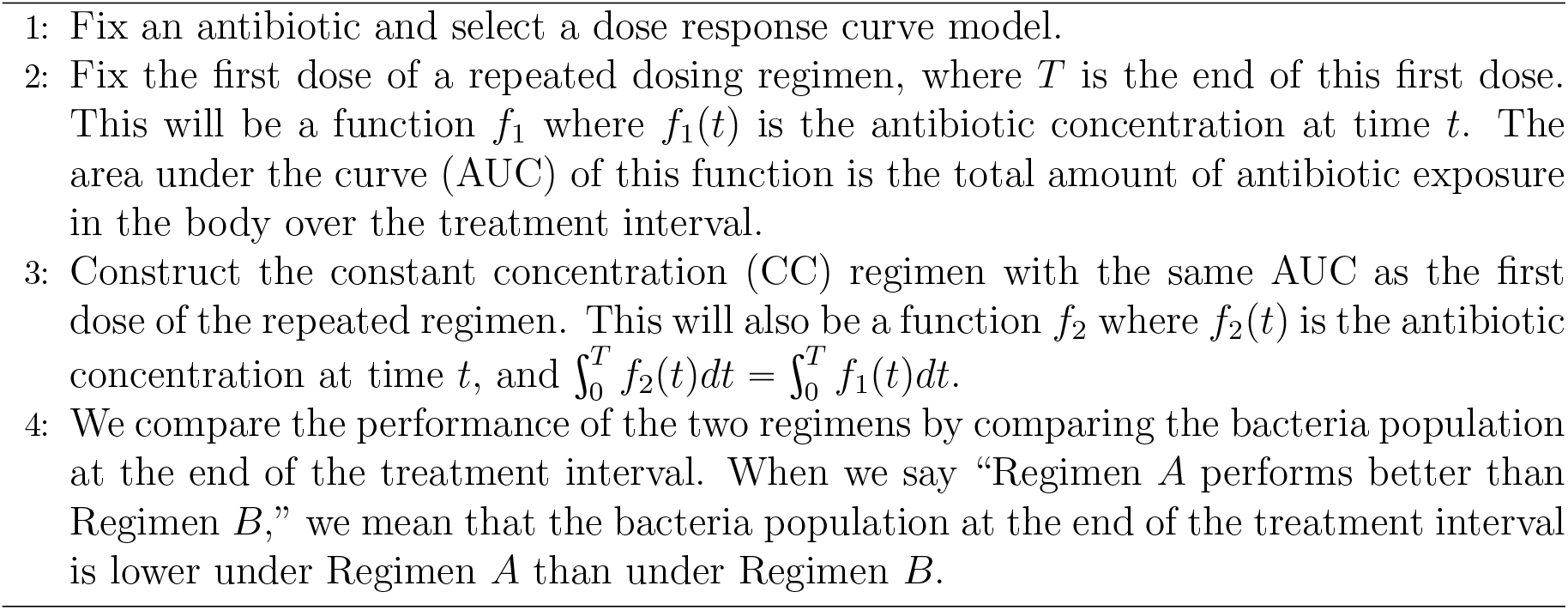

This algorithm aligns with our goal of comparing types of repeated dosing with constant concentration dosing regimens with the same total amount of antibiotic as one dose of the repeated dose regimen. Figure 1B-D illustrates three dosing regimens which satisfy our setup; they are a CC regimen and two different first doses of repeated regimens, each with the same AUC as the CC regimen.

### 2.1 Linear Dose Response Curves: All Regimens Perform Equally Well

We begin by assuming that bacterial growth is a linear function of antibiotic concentration; while less realistic than other dose response models that we will use later in this paper [2], a linear dose response curve is a good starting point for our analysis and developing the framework that we’ll continue to use throughout the paper. If the dose response is linear (i.e. if the change in net bacterial growth rate is proportional to the change in antibiotic concentration) then we will prove that all dosing regimens which satisfy our setup in Algorithm 1 will perform equally well, which will not be the case with other dose response curve shapes such as the Hill functions that we will explore later. We will then answer the question “what happens when the dose response curve is not linear?” by exploring what effects the concavity of the dose response curve have on the performance of different regimens.

We will now prove the previous claim that all dosing regimens which satisfy our setup in Algorithm 1 perform equally well under a linear dose response curve. We fix a linear dose response curve of the form

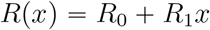

where *R*_0_, *R*_1_ ∈ ℝ and *x* ⩾ 0 is the antibiotic concentration. We fix two antibiotic regimens *f*_1_, *f*_2_ where *f*_*i*_(*t*) is the antibiotic concentration of regimen *i* at time *t*, where *f*_1_, *f*_2_ have the same AUC like in the setup in Algorithm 1; that is for a fixed *T* > 0 where *T* is the length of the treatment, we have

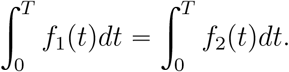

See Figure 1 for three regimens with the same AUC with this setup. Denoting the bacteria populations *b*_1_, *b*_2_ with drug concentrations *f*_1_, *f*_2_ respectively, we use the growth rate model

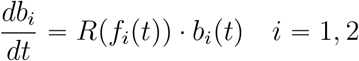

with the linear dose response curve, so our ODE becomes

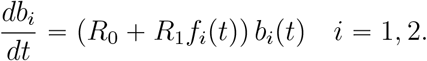

The result we are going to prove is as follows:

#### Theorem 1.

*Given a linear dose response curve R and two dosing regimens f*_1_, *f*_2_ *with the same AUC, the bacteria populations at the end of the treatment interval are equal; that is, b*_1_(*T*) = *b*_2_(*T*).

*Proof*. We define *B*_*i*_(*t*) := ln (*b*_*i*_(*t*)) so we may more easily analyze the results in log-space. We will then show *b*_1_(*T*) = *b*_2_(*T*) by proving *B*_1_(*T*) = *B*_2_(*T*) since the logarithm is a monotonically increasing function on the domain we are investigating, proving the two regimens perform equally well. Note that 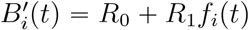. Without loss of generality, we assume *B*_1_(0) = *B*_2_(0) = 1. Then we have by basic calculus

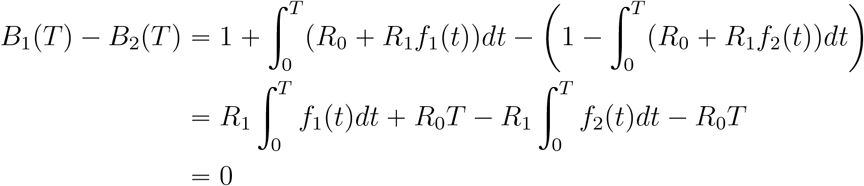

Therefore at the end of the treatment interval, both regimens have equal bacteria population. Figure 1 illustrates this result by showing the bacteria populations under the linear PD model using the three different dosing regimens with the same AUC: a constant concentration (CC) regimen and two different types of non-constant regimens that we will explore in more detail in Section 2.2.

### 2.2 “Best” Regimen Depends on the Concavity of the Dose Response Curve

Motivated mathematically by the results of the linear dose response curve, we will now explore the effects of the concavity of the dose response curve on the performance of different dosing regimens.

The effect of the antibiotic will not typically scale linearly with the drug concentration; later in this section we will be working with a standard nonlinear dose response model, a Hill function (sigmoid) [2, 15, 25]. For our analysis we consider *R* to encompass both the birth and death rates, meaning throughout the paper, *R*(*x*) = *G*_max_ − *H*(*x*) where *G*_max_ is some constant representing the maximum net growth rate in absence of antibiotics and *H*(*x*) is the Hill function describing the antibiotic-induced reduction in net growth rate. This clarification is crucial to the qualitative descriptions of our results because concavity will be important for the our analysis, and *R* will be concave down (concave) when *H* is concave up (convex), and vice versa.

#### 2.2.1 Preliminary Concavity Results

Before we explore concavity in more detail, we need to state some preliminary results that we will use later. These lemmas will aid us in proving our main concavity results, which are Theorems 2 and 3. The proofs of these lemmas can be found in Appendix B.1.

##### Lemma 1.

*Consider a function g* : [0, ∞) → ℝ.

1. *If g is strictly concave down and g*(0) = 0, *then* 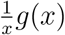 *is strictly decreasing*.
2. *If g is strictly concave up and g*(0) = 0, *then* 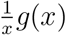 *is strictly increasing*.

##### Lemma 2.

*Given three non-colinear points* (*x*_1_, *y*_1_), (*x*_2_, *y*_2_), (*x*_3_, *y*_3_) ∈ ℝ^2^, *exactly one of the following is true:*

1. *There exists a concave up function f satisfying f continuous and f* (*x*_*i*_) = *y*_*i*_ *for i* = 1, 2, 3.
2. *There exists a concave down function f satisfying f continuous and f* (*x*_*i*_) = *y*_*i*_ *for i* = 1, 2, 3.

#### 2.2.2 Simple Pharmacokinetics: The “Step PK Model”

A common antibiotic dosing regimen is the prescription of repeated doses, such as pills or bolus injections. When repeated doses are prescribed, many pharmacokinetic processes occur such as absorption, distribution, metabolism, and excretion of the drug. These processes cause the drug concentration to vary in time, frequently resulting in decay between doses and accumulation of the drug over time. To build intuition while retaining essential features of the dynamics, we first represent repeated dosing with a simplified PK model where the antibiotic concentration is constant over a fixed interval of time [*t*_1_, *t*_2_] and zero after *t*_2_, so for now we are not considering the effects of PK processes on the drug. We will refer to this as the “step model.” One could approximately create these pulses by using an IV to keep the drug concentration constant over the dosing interval. We will spend quite a bit of time proving results about this model to build up a framework as well as an intuition for how concavity affects the efficiency of different dosing regimens. We will then extend these results to a more realistic model which does consider decay and accumulation of the drug in Section 2.3.

We will define two types of dosing regimens in this section. A constant concentration (“CC”) regimen will have a constant drug concentration *D*_1_ over the whole treatment interval [0, *T*] where *T* is the treatment length. A “pulse” regimen will have a constant drug concentration *D*_2_ over the interval [0, *T*_*p*_] and zero drug concentration over the interval [*T*_*p*_, *T*] where *T*_*p*_ is the length of the pulse. We define *AUC* to be the AUC of the CC regimen, so the CC regimen has concentration *D*_1_ = *AUC*/*T* and when we compare it to a pulse regimen of the same AUC, we have that the pulse regimen has concentration *D*_2_ = *AUC*/*T*_*p*_. The two regimens we will compare are shown in Figure 2.

We will continue to use the following bacterial growth rate model

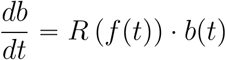

where *b* is the bacteria population (measured in any units, such as percentage of initial bacteria population), *R* is the dose-response curve and *f* is the drug concentration function. Just as we did in the proof from Section 2.1, we define *B*_*i*_(*t*) := ln (*b*_*i*_(*t*)) so we may more easily analyze the results in log-space. We denote the drug concentration function of the CC regimen as *f*_1_ and the concentration of the pulse regimen as *f*_2_. Based on our previous setup in Algorithm 1, we have *f*_1_(*t*) = *D*_1_ and

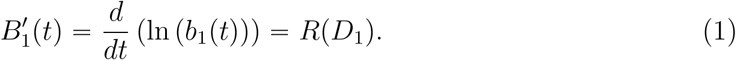

For the pulse regimen, we have 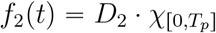 and

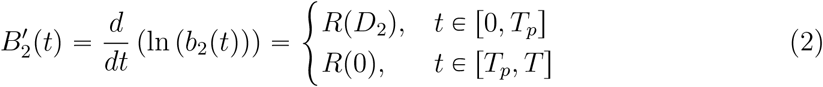

where *χ* is the indicator function. We will compare the performance of the two regimens by comparing the values of *B*_1_(*T*) and *B*_2_(*T*). Motivated by the results of Theorem 1, we want to explore the effects of the concavity of the dose response curve on the performance of the two regimens. First we will consider the case where the dose response curve is strictly concave up, i.e.

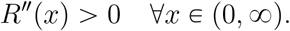

We will prove that for *R* concave up, *B*_2_(*T*) > *B*_1_(*T*), so the CC regimen performs better than the pulse regimen. We will not prove the reverse case, but we will note that the same argument with the inequalities reversed can be used to show that the CC regimen performs better than the pulse regimen when *R* is strictly concave down.

Since *R* is (strictly) concave up, we can write *R* as

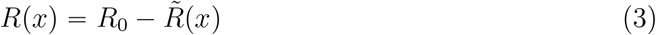

where *R*_0_ = *R*(0) and 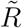 is (strictly) concave down with 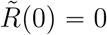. Assume *B*_1_(0) = *B*_2_(0) = *B*_0_ > 0. Then by Lemma 1, we have

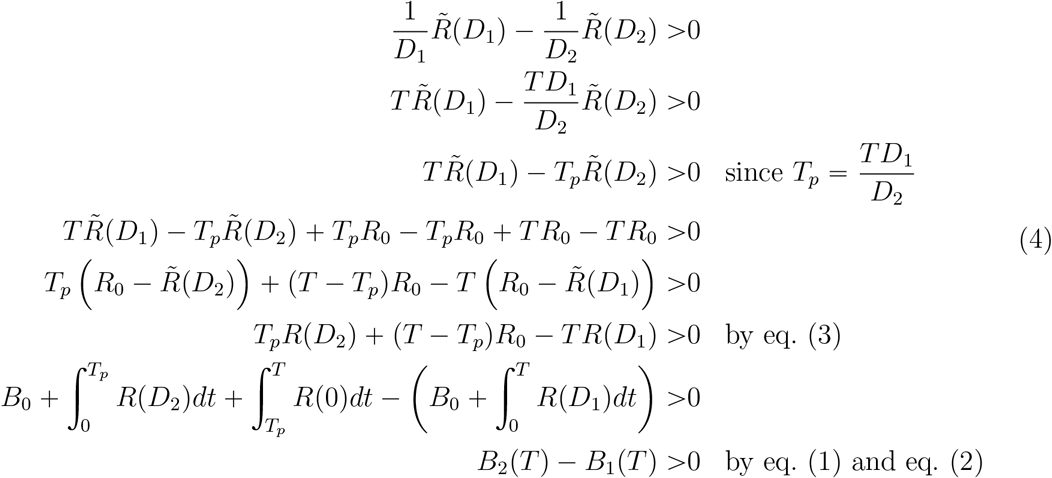

So *B*_2_(*T*) > *B*_1_(*T*), which means the CC regimen reduced the bacteria more than the pulse regimen (which we will throughout refer to as “performing better”). It is worth noting Lemma 1 uses the definition of strict concavity; if we assume non-strict concavity, we can use the same argument in (4) to show that the CC regimen performs better than or equal to the pulse regimen. We can also use (4) again but with the concavities and thus inequalities reversed to show when the dose response curve is concave up, then the pulse regimen performs better than the CC regimen.

One may then ask in the case when *R* is concave down is: Given two distinct pulse regimens, we know they will both perform better than the CC regimen, but which pulse regimen is preferred? Using (4), we see for a fixed AUC, the pulse regimen with the larger drug concentration and smaller pulse length will perform better. When *R* is concave up, the reverse is true, meaning the pulse regimen with the longer pulse length and smaller drug concentration (the regimen that more closely approximates the CC regimen with the same AUC) will perform better.

We collect the above results in the following theorem.

##### Theorem 2.

*Given a constant AUC and dose interval, the following hold true when comparing drug regimens in the Step PK Model:*

i. *If the dose-response curve is concave up, then the CC regimen performs better than any pulse regimen*.
ii. *If the dose-response curve is concave down, then any pulse regimen performs better than the CC regimen*.
iii. *If the dose response curve is concave up, then given two pulse regimens, the one with the smaller drug concentration and larger pulse length performs better*.
iv. *If the dose response curve is concave down, then given two pulse regimens, the one with the larger drug concentration and smaller pulse length performs better*.

*(When we say “Regimen A performs better than Regimen B,” we mean that the bacteria population at the end of the treatment interval is lower under Regimen A than under Regimen B.)*

Note that we may interpret any of the four parts with the word “better than” replaced with “better than or equivalent to” if we assume non-strict concavity. The distinction between strict and non-strict concavity only affects the technical steps of our proof; the results in practice are the same regardless of which convention we use to define concavity. We can see the results of this theorem applied to a particular pharmacodynamic model illustrated in Figure 6.

Note that this theorem for the Step Model holds for our already proved linear dose response curve case. Since we have noted that the results also hold for non-strict concavity by replacing “better” with “better than or equivalent to,” we observe that linear functions are both non-strictly concave up and concave down, so the CC regimen and every pulse regimen will perform equally well under a linear PD model, which agrees with what we proved in Section 2.1.

Finally, note that this result is a special case of Proposition 1 in the Appendix C, which generalizes Theorem 2 to any non-constant regimen for *A*_2_. However, we have chosen to present Theorem 2 in this main text because it is easier to understand and visualize, and it builds intuition about how convexity drives our results for the more complex results that follow when we increase both the pharmacokinetic and pharmacodynamic realism of our model.

#### 2.2.3 Adding Pharmacodynamic Realism: Hill Functions and Their Concavity

Throughout the paper so far, we have used a linear dose response curve, and we have casually referred to typical sigmoidal dose response curves, however in previous sections we have not needed to explicitly define a nonlinear dose response curve. We now look at explicit nonlinear (sigmoidal) dose response curves.

At the beginning of Section 2.2, we noted that when using a Hill function to describe the net growth rate of the bacteria as a function of antibiotic concentration, our dose response curve will be of the form *R*(*x*) = *G*_max_ − *H*(*x*) where *G*_max_ is the maximum net growth rate of the bacteria in the absence of an antibiotic and *H* is a Hill function. Whereas *R*(0) = *G*_max_ and *R* is decreasing because the effect of antibiotics is decreased net growth, *H*(0) = 0 and *H* is increasing. A common PD model which is in the form of a Hill function is the *E*_max_ model [15] whose Hill shape is dependent on the maximum efficacy of the drug (*E*_max_) and the drug concentration at which the drug has half of its maximum efficacy (*EC*_50_). A Hill function is a sigmoidal curve which can be written as

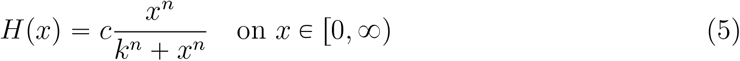

for some *n, k, c* ∈ (0, ∞). Some examples of Hill functions are shown in Figure 3. Take careful note, however, that Hill functions are more accessibly viewable on a log(*x*) scale; we later will derive the formula for the inflection points on a linear scale, which means the inflection points will appear in a different place, visually, on a log scale. The dynamics of the bacteria population under *R* are described by

**Figure 3.**
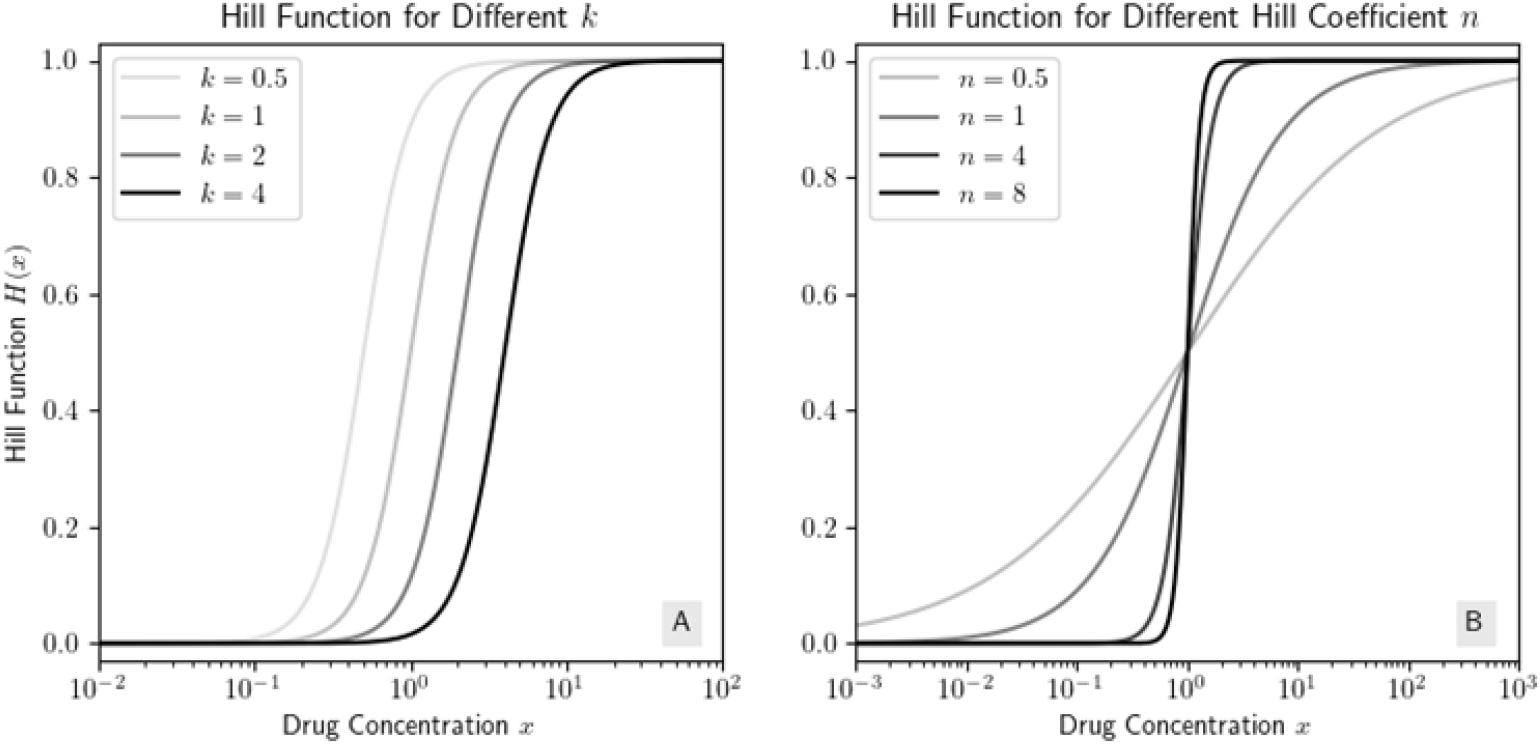
Examples of Hill functions with *c* = 1. (A) We hold *n* = 3 constant and allow *k* to vary; (B) we hold *k* = 1 constant and allow *n* to vary.

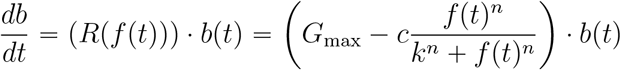

where *f* (*t*) is the drug concentration at time *t*. Note that *R*(*x*) is concave up/down if and only if *H*(*x*) is concave down/up respectively. To apply Theorem 2, we must determine the concavity of *H* by calculating its second derivative. We can rewrite *H* as

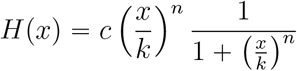

so if we set 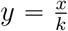 (so *y* ∈ [0, ∞)), we have

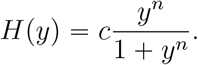

Through basic calculus (see Appendix B.2), we find

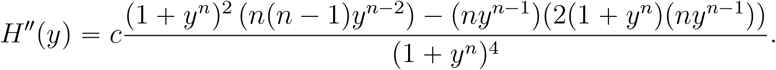

Our aim is to determine where *H* is concave up and down, so we will consider the sign of *H*″(*y*). By basic algebra, we see *H*″(*y*) = 0 if and only if (*n* − 1)*y*^*−n*^ = *n +* 1. Note 0 < *n* < 1 implies *n* − 1 < 0 but *n +* 1 > 0, so to satisfy the equation, we must have *y*^−*n*^ < 0 which is impossible for nonnegative values of *y*. So for 0 < *n* < 1, *H* has no inflection points on (0, ∞). Now, solving for *y* and assuming *n* ≠ 1, we define

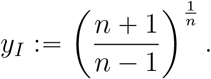

Overall we get for 0 < *n* ⩽ 1, there is no inflection point on (0, ∞), so *H* is always concave down on its domain. For *n* > 1, there is an inflection point at

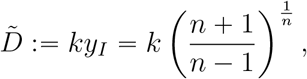

so *H* is concave up on 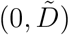 and concave down on 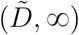.

We can now apply Theorem 2 in the following way:

I. If *n* ⩽ 1, then *H* is concave down wth *H*(0) = 0 and *R* is concave up. So for a fixed AUC and treatment interval, the CC regimen performs better than any pulse regimen.
II. If *n* > 1, then on 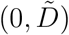, *H* is concave up with *H*(0) = 0 and *R* is concave down. So for a fixed AUC and treatment interval, if 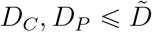 where *D*_*C*_, *D*_*P*_ are the drug concentrations of the constant and pulse regimens respectively, the pulse regimen performs better than the CC regimen.
III. If *n* > 1, then on 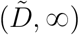, *H* is concave down. However, we do not know if we have a concave down extension of 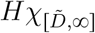 to (0, 0), so more analysis is needed.

Henceforth we will call models which are like (III) “mixed concavity models” because they have more than one concavity on their domain. Looking ahead, this case is of particular interest because the drug concentration varies during the treatment interval and may have time both above and below the inflection point. We can easily apply Theorem 2 to the leftmost concavity’s interval (like in (II)) because 0 in in the interval, so on the leftmost interval, *H* satisfies the assumptions of Lemma 1. For the step model, the points that are of interest to us are 𝒟 := {(0, 0), (*D*_*C*_, *H*(*D*_*C*_)), (*D*_*P*_, *H*(*D*_*P*_))}. In the general Hill function case, if *n* > 1 and 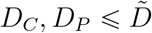, then using Lemma 2, we can draw a concave up curve Ω through 𝒟 (*H* is an example of this) and we cannot draw any concave down curve through 𝒟. So as stated, Theorem 2 readily applies.

Contrast this to (III); if *n* > 1 and 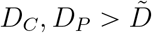, then by Lemma 2, we can either draw a concave up or concave down curve Ω through 𝒟, but not both. However, Ω’s concavity may be unrelated to *H* in that Ω may be concave up on an interval where *H* is concave down. This is illustrated in Figure 4, where Ω_1_ is concave up, even on 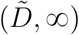 where *H* is concave down. So in order to apply Theorem 2 to any interval of the dose response curve besides the leftmost, we must first determine the concavity of Ω, which currently is defined to be just some concave up or concave down curve through 𝒟. However, by Lemma 2, we need only consider

**Figure 4.**
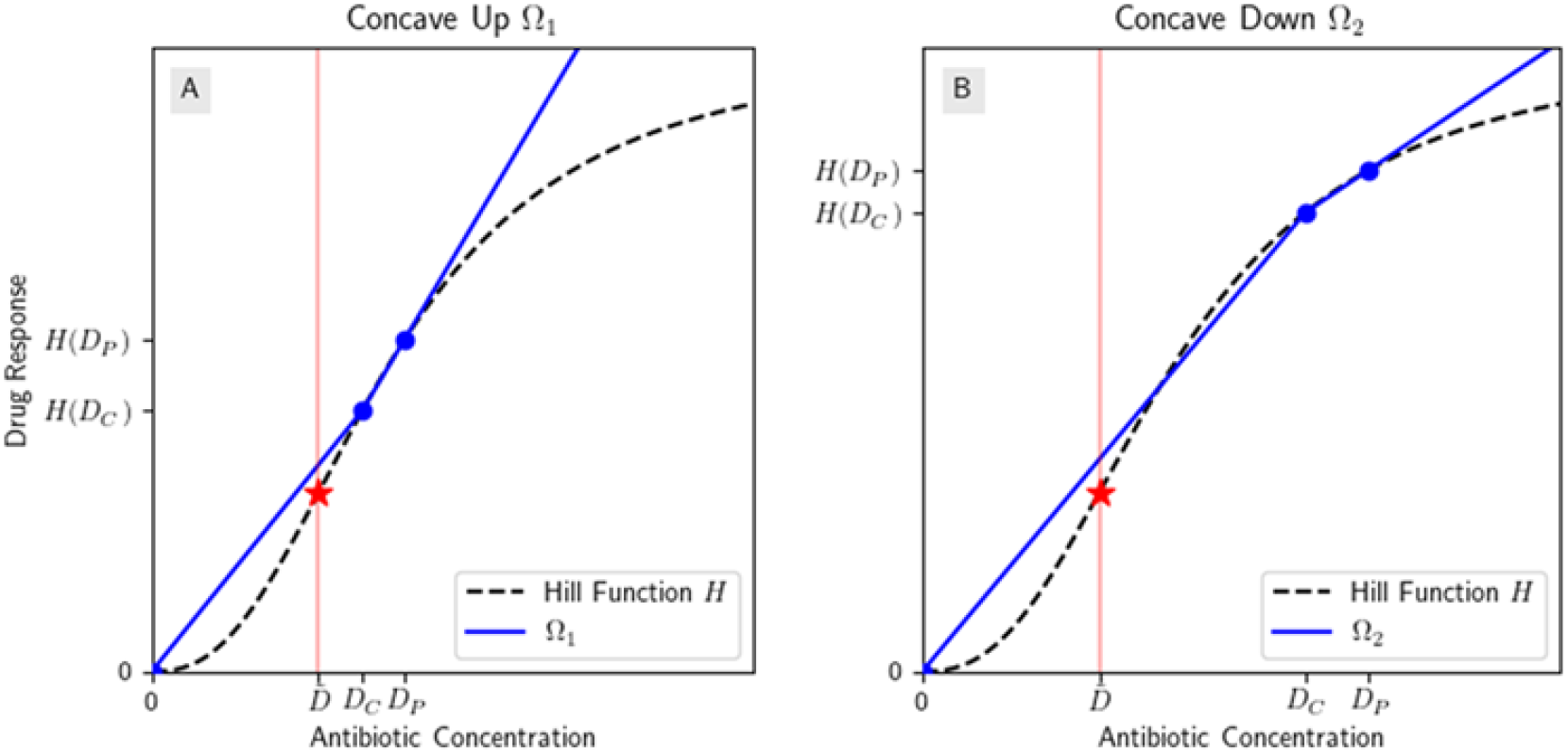
Hill functions with mixed concavities can have concave down or concave up extensions depending on {0, *D*_*C*_, *D*_*P*_}. (A) The Hill function is concave down to the right of 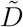, however Ω_1_ is still concave up; (B) Ω_2_ is concave down. Each graph is on a *linear* scale.

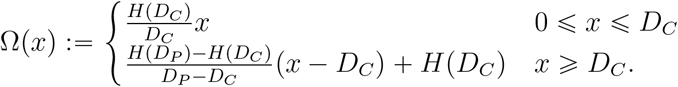

To check if Ω is concave down or up, we compare the slopes

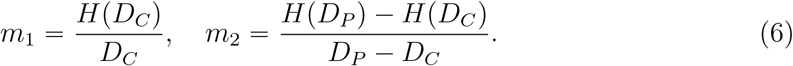

If *m*_1_ > *m*_2_, then Ω is concave down, and by Theorem 2, the CC regimen performs better than the pulse regimen. If *m*_1_ < *m*_2_, then Ω is concave up, and the pulse regimen performs better than the CC regimen.

We cannot apply Theorem 2 unless we know the concavity of Ω. Determining the concavity of Ω, or equivalently, the sign of *m*_1_ − *m*_2_, analytically for a given model may be interesting in theory, but calculating the values of *m*_1_ and *m*_2_ case-by-case may be more useful in practice because the underlying model’s complexity will not matter, and calculating *m*_1_ and *m*_2_ rely only on the values of *D*_*C*_, *D*_*P*_, *H*(*D*_*C*_), and *H*(*D*_*P*_), instead of the full model. However, we will explore this question analytically for the general Hill function as an example.

For a given *D*_*P*_, what condition must the associated *D*_*C*_ with the same AUC satisfy in order for Ω to be (WLOG) concave down? The following are equivalent to Ω being concave down:

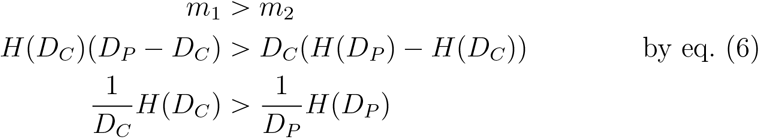

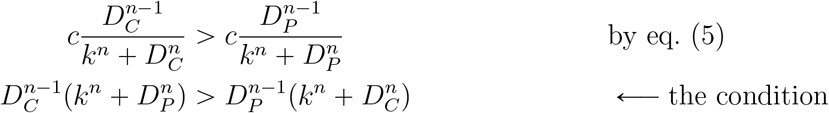

If this condition is satisfied, then *m*_1_ > *m*_2_ so Ω is concave down. If we flip all the inequalities, we have a condition for when Ω is concave up. Unfortunately in the Hill function case, these conditions are not easily solvable for *D*_*C*_ in terms of *D*_*P*_, and are more complicated than simply calculating *m*_1_ and *m*_2_. This may not always be the case for all PD models, but overall, calculating these values numerically will likely be more useful in practice.

### 2.3 Adding Pharmacokinetic Realism: The “Decay PK Model”

The step function pharmacokinetic model is an unrealistic but simple model of repeated dosing that helped us build an analytic framework. However, when pills or bolus injections are administered, decay and accumulation of the drug occur. We will refer to this model which takes first-order elimination and accumulation into account as the “decay model.” Appendix C.1 discusses how to incorporate more general pharmacokinetic models.

We will explore the decay PK model in the framework we set up for the step model and define it follows: Consider a periodic dose of concentration *A*_0_ being administered every *T* hours, where the half-life in hours of the antibiotic is *t*_half_; we then define *λ* := ln(2)/*t*_half_. Then the concentration of antibiotic *A* in the body at *t* time after one dose is given by

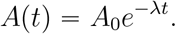

We can then recursively derive a formula for the antibiotic concentration at any time during the treatment window; we get

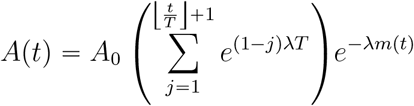

where

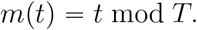

For later, note that the AUC of the first dose is 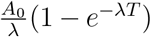, so the CC regimen with the same AUC as the first dose has drug concentration 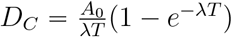 given over the interval [0, *T*]. Unlike the step model, even though identical doses are administered at equally spaced intervals so that the administration of doses is truly periodic, each “period” of the antibiotic concentration function is slightly greater than the previous period because of the accumulation of antibiotics from the previous doses, which is an aspect of dosing we did not consider in the step model. The drug concentration function of the decay model is shown in Figure 5.

**Figure 5.**
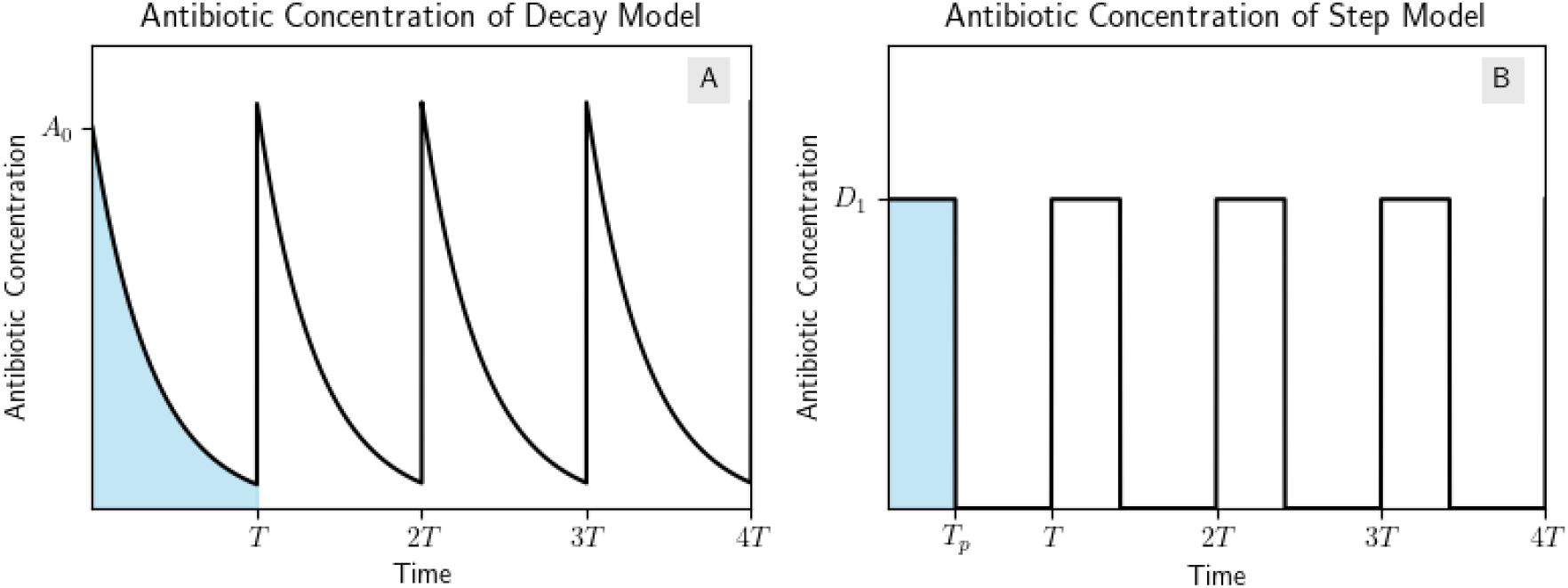
Comparing antibiotic concentration of the decay and step models. (A) The antibiotic concentration function *A*(*t*) of the decay model with initial concentration *A*_0_ and dose period *T*; (B) The antibiotic concentration function of the step model with an equal AUC as the first period of the decay model, where *T*_*p*_ is the length of the dose and *D*_1_ is the drug concentration.

**Figure 6.**
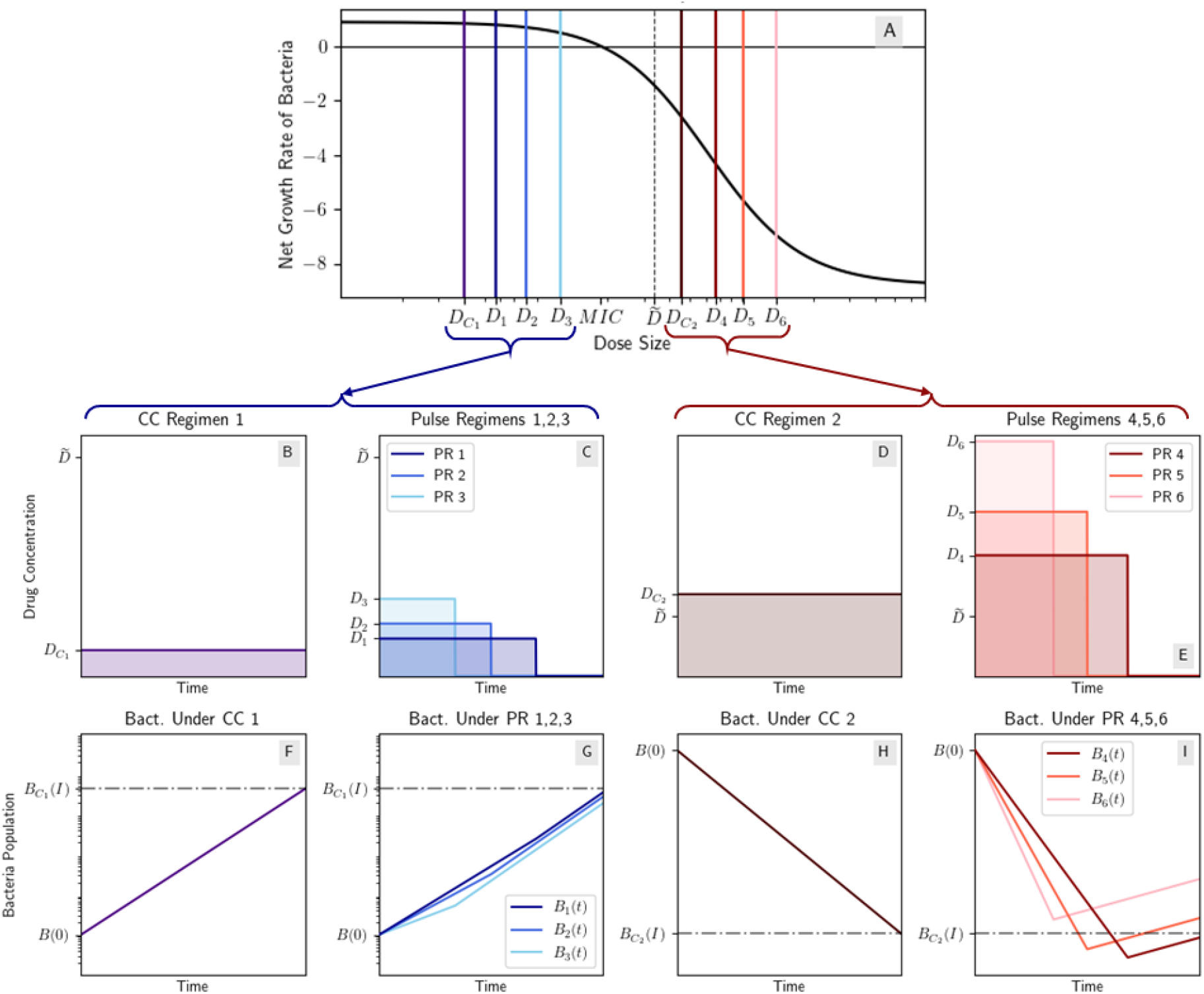
CC and pulse regimens on either side of the inflection point under a mixed concavity Hill model. Note the *y*-axes of plots on the left are scaled differently from those on the right. (A) Mixed concavity Hill dose response curve [25] where 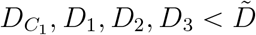 and 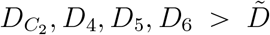; (B) CC regimen 1 with drug concentration 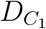; (C) three pulse regimens (PR) with the same AUC as CC 1, with drug concentrations *D*_1_, *D*_2_, *D*_3_ respectively; (D) CC regimen 2 with drug concentration 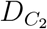; (E) three pulse regimens (PR) with the same AUC as CC 2, with drug concentrations *D*_4_, *D*_5_, *D*_6_ respectively; (F,G) the bacteria population under CC regimen 1 and PR 1,2,3; (H,I) the bacteria population under CC regimen 2 and PR 4,5,6. Note that while PR 5 and 6 perform worse than CC 2, PR 4 performs better. This is an example of the results of mixed concavity models that were discussed in Section 2.2.3. This example was constructed from data from [25] of Streptomycin being used to treat *E. coli* in vitro using *ψ*_max_ = 0.89 *h*^−1^, *ψ*_min_ = −8.8 *h*^−1^, *MIC* = 18.5 mg/L, and *n* = 1.9.

For the purposes of this section, we will study a treatment window with one periodic dose as shown in Figure 5. We also will hold constant between regimens the AUC of the drug concentration, not the total amount of antibiotic being administered (we discuss the implications of instead holding the total antibiotics constant in the Discussion). Lastly, we will assume no accumulation or exponential decay of the antibiotic in the CC regimen.

So instead of the CC regimen representing constant *administration* of an antibiotic, it could represent an IV being used to keep the concentration in the target tissue constant, corresponding to the same AUC as the periodic regimen.

Because of our choices, we can use the step model as a template for our analysis of the decay model. First we need to reprove Theorem 2 for the decay model. We do this in Appendix C, and we find that the same results hold for the decay model as for the step model. While the basic idea of the proof, taking advantage of the definition of concavity to prove our inequalities, is largely the same, the crux of this proof is in manipulating an integral to exploit concavity; in the step model, the integral was able to be resolved more easily due to the constant regions of step functions.

Just as with the proof of Theorem 2, we can replace all the strict inequalities with non-strict inequalities as well, so in the following theorem, if we assume the dose response curve is non-strictly concave down or non-strictly concave up, we may replace “better than” with “better than or equivalent to” We collect the results in the following theorem:

#### Theorem 3.

*Given a constant AUC and dose interval T, the following hold true when comparing drug regimens under the assumptions of the decay model:*

i. *If the dose response curve is concave up, then the CC regimen performs better than the periodic regimen*.
ii. *If the dose response curve is concave down, then the periodic regimen performs better than the CC regimen*.

*(When we say “Regimen A performs better than Regimen B,” we mean that the bacteria population at the end of the treatment interval is lower under Regimen A than under Regimen B.)*

*Proof*. See Appendix C.

Just as with the step model, we can see that the results of Theorem 3 agree with the results of Theorem 1 for the linear dose response curve. Also, unlike the step model, we will not compare the performance of two periodic regimens. The results would be similar to (iii) and (iv) of Theorem 2, however to construct two different periodic regimens with the same AUC, we need to alter both the dose size and the half-life of the particular drug. Unfortunately, the half-life of the drug may not be a realistic parameter to alter, so we will focus instead on a fixed periodic regimen and compare it to the CC regimen with the same AUC.

A second difference between the step and the decay model is the set of concentrations we need to consider (the range of *A*(*t*)) because this changes which parts of the dose response curve we need to analyze (the range of *R*(*A*(*t*))). In the step model, sometimes the concavity of dose response curve is not the same as the concavity of the function we draw through the discrete points (*D*_*C*_, *R*(*D*_*C*_)), (*D*_*P*_, *R*(*D*_*P*_)), and (0, 0). Intuitively, it is the addition of the point (0, 0) that causes this discrepancy. In the decay model, however, we consider the entire connected interval of doses between the minimum and (*D*_min_) maximum (*D*_max_) drug concentrations of the periodic regimen, so we do not have this issue. The results of Theorem 3 may be rewritten in terms of the concavity *R* solely on the interval of concentrations attained by the decay periodic regimen. We can replace “If the dose response curve is concave up/down” with “If the dose response curve is concave up/down on the interval (*D*_min_, *D*_max_)” and indeed the proof provided in Appendix C generalizes to a connected interval of doses instead of the entire domain of *R*. If one wishes to confirm this intuition explicitly akin to Figure 4, see Appendix D.

Throughout our analytic work, we have established that the concavity of the dose response curve is important to determining which regimen performs better. This analysis rests on shape of the full dose response curve and not just one single parameter such as the *EC*_50_ or the MIC. When a dose response curve *R* is concave up, the function describing the reduction in the net growth rate of the bacteria (what we called *H* for Hill functions) is concave down, which means the phenomenon of diminishing returns informs the performance of the regimens. In this case, CC regimens perform better because the associated periodic regimen must start at a higher drug concentration to achieve the same AUC.

## 3 Application of Results to Real Antibiotic Dosing

In this section we demonstrate our theoretical results with real, empirically obtained antibiotic parameters. We will demonstrate our results first with the step PK model as a toy example for illustrative purposes, and then the decay PK model as the more realistic example with real antibiotic dosing regimens considered. For these tasks, we will use the Hill function PD model as given in Regoes et al. [25], which is a reparametrization of the standard *E*_max_ model [2, 15], and is given by

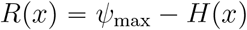

where [25] defines their Hill function *H* as

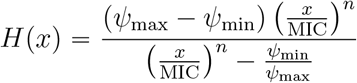

- *ψ*_max_ is the maximum net growth rate of the bacteria in absence of an antibiotic (which we called *G*_max_ earlier).
- *ψ*_min_ is the minimum net growth rate of the bacteria in the presence of an antibiotic. Note when *ψ*_min_ < 0, this indicates that at high antibiotic concentration, the bacteria population is decaying.
- MIC is the minimum inhibitory concentration of the antibiotic, which is the concentration where the net growth rate *R*(MIC) = 0. Regoes et al. estimated the MIC both using twofold dilution and by fitting a Hill function to time-kill curve data [25]; we use their fitted values.
- *n* is the Hill coefficient

We can rewrite *H* as

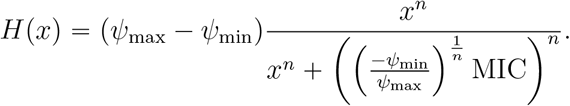

To put this equation into the general form of a Hill function, we set *c* = (*ψ*_max_ − *ψ*_min_) and 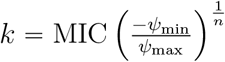. Then if *n* > 1, our inflection point occurs at

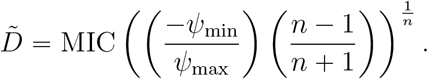

To demonstrate our results with the step model, we apply Theorem 2. The following list of results hold true for this reparametrized Hill model:

1. If *n* ⩽ 1, *H* is concave down, so the CC regimen performs better than any pulse regimen.
2. If *n* > 1 and 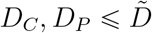, then the pulse regimen performs better than the CC regimen.
3. If *n* > 1 and 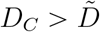 or 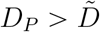, then we must determine the concavity of Ω as discussed at the end of Section 2.2.3 to determine which regimen performs better.

Figure 6 illustrates cases (II) and (III) using toy dosing regimens with streptomycin parameters from [25]. Streptomycin was discovered in 1944 and was the first antibiotic used to treat *M. tuberculosis* [9].

To numerically demonstrate our results with the decay model, we first note that the list of results from the end of Section 2.3 will hold true for this parametrization as well by setting

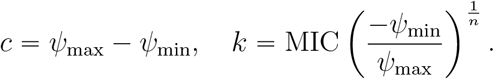

In the following, we will first consider ampicillin with the parametrized model and parameters from [25]. In that study, dose-response curves of various antibiotics’ effect on the net growth rate of *E. coli* were fit to collected data. We define a few periodic regimens and their associated CC regimens and simulate the bacteria population over the treatment window. We will compare the performance of the periodic regimens with their associated CC regimens (note for ampicillin, since *n* = 0.75 < 1, each CC regimen will perform better). We will then compare and contrast ampicillin’s results with rifampin and ciprofloxacin. We chose these antibiotics from [25] to compare and contrast the results when the Hill coefficient *n* < 1 (ampicillin, *n* = 0.75) with different cases when *n* > 1 (rifampin, *n* = 2.5 and ciprofloxacin, *n* = 1.1).

The code used to generate the numerical results in this paper is available at https://github.com/leahchilders/GeneralAntibioticDosing.

### 3.1 Numerical Results When Hill Coefficient c 1: The Case of Ampicillin

Ampicillin is a primarily bactericidal *β*−lactam antibiotic used to treat a number of bacterial infections [5] such as *Streptococcus pneumoniae* [35] and is an example of an antibiotic whose fitted dose response curve from [25] had its Hill coefficient *n* < 1. Table 1 contains the other PK and PD parameters from [25] we use to illustrate the regimen performance of ampicillin.

**Table 1:** Three ampicillin regimens. Constant concentration is calculated as the AUC of the periodic dose divided by the dose period. Half-life from [29], volume of distribution from [11], PD parameters from [25].

| Parameter | Regimen 1 | Regimen 2 | Regimen 3 |
| --- | --- | --- | --- |
| Half-life (h) | 1.7 |  |  |
| Volume of distribution (L) | 17.29 |  |  |
| Periodic dose size (mg) | 250 | 500 | 1000 |
| Periodic initial conc. (mg/L) | 14.46 | 28.92 | 57.84 |
| Dose period (h) | 6 | 4 | 8 |
| $AUC$ (mg · h/L) | 32.39 | 57.04 | 136.41 |
| Constant conc. (mg/L) | 5.40 | 14.26 | 17.05 |
| $\psi_{\max}$ (h<sup>-1</sup>) | 0.75 | | |
| $\psi_{\min}$ (h<sup>-1</sup>) | −4.0 | | |
| MIC (mg/L) | 3.4 |  |  |
| $n$ | 0.75 | | |

We give three different periodic regimens as examples. Using an average human weight of 70 kg we get from [11] a volume of distribution of Ampicillin of ≈ 17.29 L. Regimen 1 has a dose size of 250 mg and a dose period of 6 hours. Thus the initial drug concentration is 50 mg/L and the constant concentration (CC) regimen with the same AUC (32.29 mg/L) has concentration 5.40 mg/L · h. This means we are administering ampicillin at a rate around 93 mg/h. These values are also calculated for regimens 2 and 3, and contained in Table 1.

Since the Hill coefficient *n* < 1, the dose response curve for ampicillin is always concave down, so for each periodic regimen, its respective CC regimen will perform better than the periodic regimen. The dose response curve, drug concentration functions, and bacteria population curves or each regimen are shown in Figure 7.

**Figure 7.**
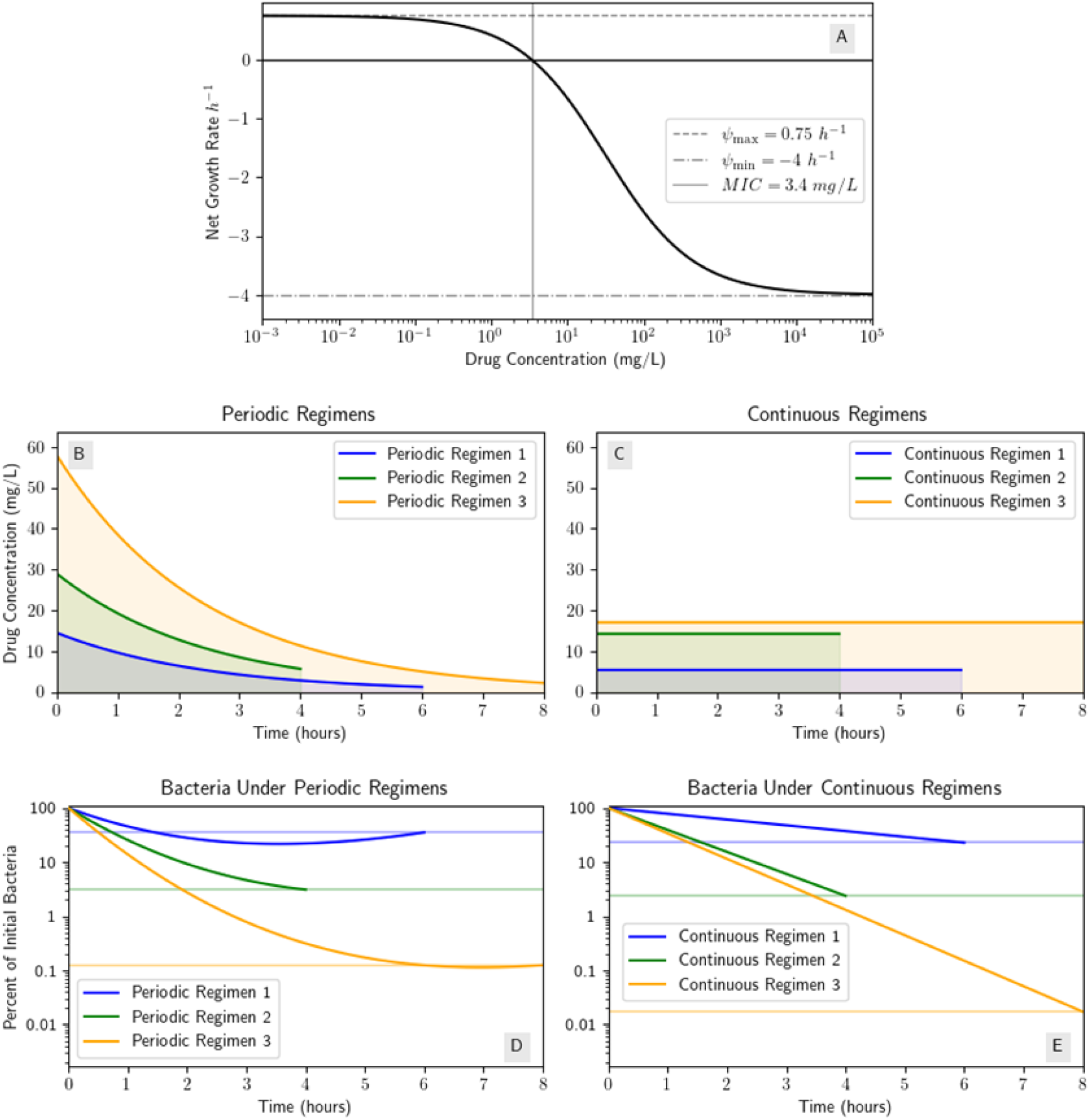
Numerical simulations for ampicillin and various dosing regimens demonstrate the CC regimen always performs better because *n* < 1. (A) Dose response curve for ampicillin [25]; (B) periodic dosing regimens described in Table 1; (C) the periodic regimens’ respective CC regimens (i.e. CC regimen *i* has the same AUC as periodic regimen *i*); (D) the bacteria population under each periodic regimen; (E) the bacteria population under each CC regimen.

Tetracycline, one of the other antibiotics used in [25] and used to treat many bacterial infections such as Lyme disease [7], is another example of an antibiotic with *n* < 1 and the results for tetracycline would be the same as the results for ampicillin; each periodic regimen will perform worse than its associated CC regimen. Thus, for both ampicillin and tetracycline, dosing with a periodic regimen may not reduce the bacteria population at the end of the treatment as much as the constant concentration regimen with the same AUC or antibiotic exposure.

### 3.2 Numerical Results When Hill Coefficient > 1: The Case of Ciprofloxacin and Rifampin

Ciprofloxacin is a broad-spectrum antibiotic used to treat many bacterial infections including *Bacillus anthracis* [30] whose dose response curve has Hill coefficient *n* = 1.1 > 1 [25], so unlike ampicillin, ciprofloxacin’s dose response curve is of mixed concavity. Also of mixed concavity are rifampin, used to treat tuberculosis and whose dose response curve has *n* = 2.5 > 1, and streptomycin, whose dose response curve has *n* = 1.9 > 1 [25]. We will compare ciprofloxacin and rifampin to ampicillin. Ciprofloxacin’s Hill coefficient is close to 1, so the inflection point is comparatively close to 0; on the other hand rifampin’s Hill coefficient is much larger, so the inflection point is much larger. As we will see from the results of both of the antibiotics, in order to define the regimens we want to compare for each antibiotic, we must choose very small doses for cirpofloxacin and very large doses for rifampin. This will have implications of the realism of the regimens we define, meaning some of the regimens we define will not be accessible in practice. Figures E2 in Appendix E.1 show the dose response curves of ciprofloxacin and rifampin, as well as streptomycin and tetracycline.

Using the values given in Table E2 in the Appendix E.2, we get that ciprofloxacin’s inflection point 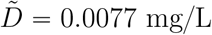, so we have three possible cases: (i) if our interval of concentrations 𝒟 = (*D*_min_, *D*_max_) is less than 0.0077, the periodic regimen will perform better; (ii) if 𝒟 is greater than 0.0077, the CC regimen will perform better; (iii) if *D*_min_ < 0.0077 < *D*_max_, we cannot analytically determine which regimen will perform better and we must examine the numerical result. Table 2 gives four periodic regimens and both the periodic and associated CC regimens’ end bacteria populations. Regimen 2 is an example of case (i), regimen 1 is an example of case (ii), and regimens 3 and 4 are examples of case (iii). Appendix E.2 contains all the details of the four dosing regimens; Table 2 gives the dose sizes, treatment lengths, and end bacteria populations, and Figure E3 shows the bacteria population curves for each regimen.

**Table 2:** Four ciprofloxacin regimens and the bacteria population value at the end of each treatment interval. End bacteria population values are given as a percentage of the initial bacteria population.

| Parameter | Regimen 1 | Regimen 2 | Regimen 3 | Regimen 4 |
| --- | --- | --- | --- | --- |
| Periodic dose size (mg) | 100 | 1 | 30 | 2.5 |
| Dose period (h) | 6 | 12 | 24 | 12 |
| Periodic bac. end value | $4.46 \times 10^{-10}$ | $1.48 \times 10^6$ | $1.97 \times 10^{-2}$ | $3.00 \times 10^5$ |
| Constant bac. end value | $2.34 \times 10^{-10}$ | $1.50 \times 10^6$ | $1.67 \times 10^{-4}$ | $3.04 \times 10^5$ |
| Which regimen performed better? | Continuous | Periodic | Continuous | Periodic |

Using the values given in Table E3, we get that rifampin’s inflection point 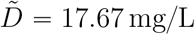, so similarly to ciprofloxacin, we have the same three possible cases: (i) if our interval of concentrations 𝒟 = (*D*_min_, *D*_max_) is less than 17.67, the periodic regimen will perform better; (ii) if 𝒟 is greater than 17.67, the CC regimen will perform better; (iii) if *D*_min_ < 17.67 < *D*_max_, we cannot analytically determine which regimen will perform better and we must examine the numerical result. Table 3 gives five periodic regimens with varying dose sizes and compares the regimens’ performance to the performance of the associated CC regimen. Contrasting ciprofloxacin, the concave up domain of rifampin’s dose response curve (so the concave down domain of the Hill function) is practically inaccessible. Regimen 5 is an example of case (i), regimens 2, 3, and 4 are examples of case (ii), and regimen 1 is an example of case (iii). Appendix E.2 contains all the details of the five dosing regimens; Table 3 gives the dose sizes, treatment lengths, and end bacteria populations, and Figure E4 shows the bacteria population curves. For the half-life value, while it is known that the half-life of rifampin can change based on the treatment regimen [37], we will hold it constant at *t*_half_ = 2.5 hours. However, using the provided code, one may test other values of the half-life if desired.

**Table 3:** Five rifampin regimens and the bacteria population value at the end of each treatment interval. End bacteria values are given as a percentage of the initial bacteria population.

| Parameter | Reg. 1 | Reg. 2 | Reg. 3 | Reg. 4 | Reg. 5 |
| --- | --- | --- | --- | --- | --- |
| Per. dose size (mg) | 7000 | 3100 | 3000 | 1400 | 500 |
| Dose period (h) | 6 | 12 | 12 | 12 | 12 |
| Per. end bac. value | $7.93 \times 10^{-8}$ | $4.12 \times 10^{-2}$ | $6.98 \times 10^{-2}$ | $1.73 \times 10^3$ | $2.41 \times 10^5$ |
| Cont. end bac. value | $8.04 \times 10^{-9}$ | $2.76 \times 10^{-2}$ | $7.25 \times 10^{-2}$ | $2.24 \times 10^4$ | $3.50 \times 10^5$ |
| Which reg. performed better? | Continuous | Continuous | Periodic | Periodic | Periodic |

The data from [25] for the antibiotics we have been using throughout this section is from in vitro studies of *E. coli*, which is not a bacteria rifampin is typically used to treat, but nicely illustrates some important clinical implications of our results. Recent studies have shown that doses up to 35-40 mg/kg daily of rifampin are well-tolerated [4, 27, 33] but doses of 50 mg/kg daily are not [33]. For our average 70 kg human, 40 mg/kg and 50 mg/kg would be 2800 mg and 3500 mg respectively. As shown in Table 3, these dose sizes are near the threshold above which the CC regimen performs better than the periodic regimen. This issue is compounded by the half-life of rifampim being around 2.5 hours [37] and the dose periods we have defined being 6-12 hours, meaning the AUC of these regimens will be around 10,000-20,000 mg · h/L. This means the associated CC regimens will have to administer rifampin (with an IV, so bioavailability is not a concern) at 20,000-40,000 mg/day, which is even more drug per day than the daily 50 mg/kg dose that is known to not be well-tolerated. In other words, both the periodic and CC dosing regimens that would be necessary to yield a better performance for the CC regimen are not realistic. In the nontoxic doses ranges of rifampin against *E. coli*, the periodic regimen will always perform better than the CC regimen.

These results for rifampin contrast ciprofloxacin, where realistic dosing regimens will be in the domain of the dose response curve where the associated CC regimen performs better than the periodic regimen, even though both antibiotics have mixed concavity dose response curves.

Overall, our results suggest that there is not a universal best dosing regimen for antibiotics. Whether repeated or constant doses are better depends on the antibiotic properties and the desired or nontoxic dosing range. In particular for our work, the better strategy depends on the shape of the entire dose response curve and not just one single PK/PD parameter value.

## 4 Discussion

In this paper, we proved analytic results about choices of dosing regimens based on the concavity of dose response curves in the context of a standard bacterial growth rate model, which we then applied to different dose response curves; in particular, we explored linear dose response curves and Hill functions. We identified (analytically when possible, numerical when necessary) a candidate for the best regimen dependent on the antibiotic’s PD/PK parameters (half-life, *E*_*max*_, *EC*_50_, MIC, etc) and the repeated treatment regimen’s parameters (treatment length, initial drug concentration, etc).

The results of our exploration suggest that the maxim “hit hard and hit early,” which is starting treatment with a high antibiotic dose immediately to rapidly reduce the bacterial load, may not always be the best strategy to optimize antibiotic treatment. For example, in the case of ampicillin whose Hill coefficient *n* = 0.75 < 1, our analysis claims a lower-dosed constant concentration regimen may perform better (i.e. the bacteria population will be lower at the end of the treatment interval) than a repeated regimen in either the step or decay model. This means that if we are able to keep the drug concentration of all regimens above the MIC of the drug, then even though all regimens will eventually witness the death of the bacteria population, we will reduce the population faster by administering the antibiotic at a constant rate. This is in contrast to the FDA recommendation of dosing IV ampicillin periodically [35].

Here, we used a very simple model utilizing only the dose response curve of the antibiotic for the bacteria as a single population. However, there exist more realistic and complex models describing the pharmacodynamics of the antibiotic. One such example is the drug-target binding model described in [13] which takes into account interactions between the antibiotic and the bacteria subpopulations on a molecular level. The introduction of subpopulations of bacteria with varying susceptibilities to the antibiotic as well as the added complexity of drug-target binding kinetics would likely change the results of our analysis to be more nuanced but also more realistic. Performing similar analysis on these more complex models would be a very interesting extension of this work which may yield more nuanced and complicated results, still dependent on the full profile of the PD model.

In deriving our results, we necessarily made assumption choices, such as the choice that we are holding the AUC of the drug concentration function of each regimen constant between the regimens we compare. This means that the total amount of antibiotic administered in each regimen may be different, especially when we consider accumulation and decay of the drug. This choice was intentional as the AUC represents the total antibiotic exposure over time. However, optimizing the dosing regimen holding constant the total amount of antibiotic used may also be desired, such as when dealing with a limited drug supply. Decades of work has been done on this question as well [8], and a natural extension of our work would be to explore the query “given a fixed *X*, what dosing regimen minimizes the bacteria population at the end of the treatment window?” where *X* could be any measured quantity we wish to hold constant between regimens.

We also neglected the possibility of drug resistance, a key concern in the field of antibiotic dosing. We assumed the bacteria population was composed homogeneously of bacteria which always responds to the antibiotic in the same way as the rest of the bacteria in the population. However, we know this is not always true, and a bacteria population is a heterogeneous population of different strains with varying levels of susceptibility, thus varying levels of response to the drug [1]. The purpose of the present study was to develop the theoretical framework to explore questions about comparing dosing regimens; in the future, similar questions should be explored with models (such as the 2-dimensional ODE model from [34]) which account for the emergence of drug resistant strains.

However, neglecting resistance also means that, in our framework, we do not account for mutant selection, specifically the minimal selective concentration (MSC), which occurs sub-MIC where *de novo* mutants may be selected for [12], and the mutant selection window (MSW), a range of concentrations between the MIC and the mutant prevention concentration (MPC) where first-step mutants may be enriched. Dosing regimens which keep the drug concentration above the MSW are typically preferred to avoid the emergence of resistance, while our framework only considers minimizing the total bacteria population at the end of the treatment length. In these cases, given a fixed antibiotic supply, it would be necessary to evaluate the tradeoff between the two goals. These concerns can be addressed in future work using a model which explicitly accounts for resistant strains and their emergence, where one can evaluate both the total bacteria population and the probability of resistance emergence under different dosing regimens.

Past studies have explored similar questions to ours in an experimental setting for very specific cases [10, 20, 36], however we ask this question in an analytical setting to explore a wider range of cases. While some papers have noted the ceiling effect caused by the concave down shape of Hill functions [2, 14], we build on this by exploring the best dosing strategy based on the shape of the entire dose response curve, including in what we have named “mixed concavity models.” Our results also complement other modeling results such as those in [17] who analytically determined the dosing regimen which minimizes the AUC of the drug concentration function. The author of [17] also determined, given a fixed AUC, the best “constant” regimen to minimize the bacterial load. However, their definition of “constant” regimen is different from ours; they define a constant regimen as one where the drug concentration is held constant for a certain amount of time that might be less than *T*; i.e. their variable *T*_opt_ is the equivalent of our variable *T*_*P*_, the length of the dose in the periodic regimen under the Step PK model, *not* our variable *T*, the dose period, and in our framework, essentially the treatment window. They do not allow the bacteria population to regrow following the end of the dose, instead measuring the bacterial load at the end of the dose, not the treatment window. Allowing *T* to be a free variable or fixing *T* describe different practical situations and considerations. One may prefer allowing *T* to be a fixed as we did when one wants to extend the treatment to be periodic over a longer time frame than beyond the initial dose, because the period of regrowth impacts the success of the overall treatment as opposed to optimizing for the minimum bacterial load at the end of a single dose.

Furthermore, our results parallel the concept of “antifragility” applied by Taleb and West to risk management in finance and oncology [32]. They find that systems with concave up response curves benefit from variability whereas concave down systems suffer under variability. This is directly in line with our results: when the magnitude of bacterial dose response 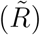 is concave down, the treatment “suffers” with higher variability (antibiotic concentration fluctuations), and when the magnitude of response is concave up, the treatment benefits with fluctuating antibiotic concentration.

Our results add to the growing body of literature suggesting that there is no one-size-fits-all dosing regimen for antibiotic treatments, challenging the use of “hit hard and hit early” as the universal rule. The choice between high-dose repeated regimens or low-dose constant regimens depends on the specific antibiotic, the target dosing range, and most notably in our work, the overall shape of the dose-response curve, rather than relying on a single PK/PD parameter value.

## Acknowledgements

JMC acknowledges the support of the National Institutes of Health (grant nos. R21-AI143443-01A1 and R01-OD011095). PAzW acknowledges the support of the Norwegian Research Council (grant no. 262686).

The authors declare the following financial interests/personal relationships which may be considered as potential competing interests: JMC has served as a consultant for Excision BioTherapeutics and Merck.

We thank Lilith Loriinae, Dennis Partl, and Jingyi Liang for their helpful comments on the basic ideas of this paper. We also thank Jonathan Jenkins for his analytical insights on the implications of Theorem 2.

## References

[1] Asgher Ali, Mudassar Imran, Sultan Sial, and Adnan Khan. Effective antibiotic dosing in the presence of resistant strains. PLoS One, 17(10):e0275762, 2022.

[2] Gunnar Alván, Gilles Paintaud, and Monique Wakelkamp. The efficiency concept in pharmacodynamics. Clinical Pharmacokinetics, 36:375–389, 1999.

[3] Kimberly G Blumenthal, Jonny G Peter, Jason A Trubiano, and Elizabeth J Phillips. Antibiotic allergy. The Lancet, 393(10167):183–198, 2019.

[4] Martin J Boeree, Andreas H Diacon, Rodney Dawson, Kim Narunsky, Jeannine Du Bois, Amour Venter, Patrick PJ Phillips, Stephen H Gillespie, Timothy D McHugh, Michael Hoelscher, et al. A dose-ranging trial to optimize the dose of rifampin in the treatment of tuberculosis. American Journal of Respiratory and Critical Care Medicine, 191(9):1058–1065, 2015.

[5] Karen Bush and Patricia A Bradford. β-lactams and β-lactamase inhibitors: an overview. Cold Spring Harbor Perspectives in Medicine, 6(8):a025247, 2016.

[6] Jean Carlet, Peter Collignon, Don Goldmann, Herman Goossens, Inge C Gyssens, Stephan Harbarth, Vincent Jarlier, Stuart B Levy, Babacar N’Doye, Didier Pittet, et al. Society’s failure to protect a precious resource: antibiotics. The Lancet, 378(9788):369–371, 2011.

[7] Ian Chopra and Marilyn Roberts. Tetracycline antibiotics: mode of action, applications, molecular biology, and epidemiology of bacterial resistance. Microbiology and Molecular Biology Reviews, 65(2):232–260, 2001.

[8] Hubert C Chua and Vincent H Tam. Optimizing clinical outcomes through rational dosing strategies: roles of pharmacokinetic/pharmacodynamic modeling tools. In Open Forum Infectious Diseases, volume 9, page ofac626. Oxford University Press US, 2022.

[9] Keira A. Cohen, Katharine E. Stott, Vanisha Munsamy, Abigail L. Manson, Ashlee M. Earl, and Alexander S. Pym. Evidence for expanding the role of Streptomycin in the management of drug-resistant Mycobacterium tuberculosis. Antimicrobial Agents and Chemotherapy, 64(9):10.1128/aac.00860–20, 2020.

[10] WA Craig and SC Ebert. Continuous infusion of beta-lactam antibiotics. Antimicrobial Agents and Chemotherapy, 36(12):2577–2583, 1992.

[11] Mats Ehrnebo, Sten-Ove Nilsson, and Lars O Boréus. Pharmacokinetics of ampicillin and its prodrugs bacampicillin and pivampicillin in man. Journal of Pharmacokinetics and Biopharmaceutics, 7(5):429–451, 1979.

[12] Erik Gullberg, Sha Cao, Otto G Berg, Carolina Ilbäck, Linus Sandegren, Diarmaid Hughes, and Dan I Andersson. Selection of resistant bacteria at very low antibiotic concentrations. PLoS pathogens, 7(7):e1002158, 2011.

[13] Colin Hemez, Fabrizio Clarelli, Adam C Palmer, Christina Bleis, Sören Abel, Leonid Chindelevitch, Theodore Cohen, and Pia Abel Zur Wiesch. Mechanisms of antibiotic action shape the fitness landscapes of resistance mutations. Computational and Structural Biotechnology Journal, 20:4688–4703, 2022.

[14] Nick Holford. Pharmacodynamic principles and the time course of immediate drug effects. Translational and Clinical Pharmacology, 25(4):157–161, 2017.

[15] Nick Holford and Lewis Sheiner. Understanding the Dose-Effect Relationship. Clinical Pharmacokinetics, 6, 12 1981.

[16] Edward W Hook, Kimberly Workowski, Jodie A Dionne, Candice J McNeil, Stephanie N Taylor, Batteiger Teresa, Julia C Dombrowski, Kenneth H Mayer, Arlene C Seña, Harold C Wiesenfeld, Charlotte Perlowski, Lori Newman, Chunming Zhu, and Jorge E Mejia-Galvis. One vs Three Weekly Doses of Benzathine Penicillin G for Treatment of Early Syphilis in Persons with and without HIV: A Multicenter Randomized Controlled Trial (RCT). Open Forum Infectious Diseases, 10, 11 2023.

[17] Guy Katriel. Optimizing antimicrobial treatment schedules: some fundamental analytical results. Bulletin of Mathematical Biology, 86(1):1, 2024.

[18] Ping Liu, Markus Müller, and Hartmut Derendorf. Rational dosing of antibiotics: the use of plasma concentrations versus tissue concentrations. International Journal of Antimicrobial Agents, 19(4):285–290, 2002.

[19] Carl Llor and Lars Bjerrum. Antimicrobial resistance: Risk associated with antibiotic overuse and initiatives to reduce the problem. Therapeutic Advances in Drug Safety, 5:229–241, 12 2014.

[20] Kirsten J. Meyer, Hannah B. Taylor, Jazlyn Seidel, Michael F. Gates, and Kim Lewis. Pulse Dosing of Antibiotic Enhances Killing of a Staphylococcus aureus Biofilm. Frontiers in Microbiology, 11, 2020.

[21] Samiha Mohsen, James A Dickinson, and Ranjani Somayaji. Update on the adverse effects of antimicrobial therapies in community practice. Canadian Family Physician, 66(9):651–659, 2020.

[22] Markus Mueller, Amparo de la Pena, and Hartmut Derendorf. Issues in pharmacokinetics and pharmacodynamics of anti-infective agents: kill curves versus MIC. Antimicrobial Agents and Chemotherapy, 48(2):369–377, 2004.

[23] Elisabet I Nielsen, Otto Cars, and Lena E Friberg. Pharmacokinetic/pharmacodynamic (PK/PD) indices of antibiotics predicted by a semimechanistic PKPD model: a step toward model-based dose optimization. Antimicrobial Agents and Chemotherapy, 55(10):4619–4630, 2011.

[24] Paolo S Ocampo, Viktória Lázár, Balázs Papp, Markus Arnoldini, Pia Abel zur Wiesch, Róbert Busa-Fekete, Gergely Fekete, Csaba Pál, Martin Ackermann, and Sebastian Bonhoeffer. Antagonism between bacteriostatic and bactericidal antibiotics is prevalent. Antimicrobial Agents and Chemotherapy, 58(8):4573–4582, 2014.

[25] Roland Regoes, Camilla Wiuff, Renata Zappala, Kim Garner, Fernando Baquero, and Bruce Levin. Pharmacodynamic Functions: a Multiparameter Approach to the Design of Antibiotic Treatment Regimens. Antimicrobial Agents and Chemotherapy, 48:3670–6, 10 2004.

[26] Rovina Ruslami, Hanneke MJ Nijland, Bachti Alisjahbana, Ida Parwati, Reinout van Crevel, and Rob E Aarnoutse. Pharmacokinetics and tolerability of a higher rifampin dose versus the standard dose in pulmonary tuberculosis patients. Antimicrobial Agents and Chemotherapy, 51(7):2546–2551, 2007.

[27] Charlotte Seijger, Wouter Hoefsloot, Inge Bergsma-de Guchteneire, Lindsey Te Brake, Jakko van Ingen, Saskia Kuipers, Reinout Van Crevel, Rob Aarnoutse, Martin Boeree, and Cecile Magis-Escurra. High-dose rifampicin in tuberculosis: experiences from a Dutch tuberculosis centre. PLoS One, 14(3):e0213718, 2019.

[28] Garima Singh, Mehmet A Orman, Jacinta C Conrad, and Michael Nikolaou. Systematic design of pulse dosing to eradicate persister bacteria. PLoS Computational Biology, 19(1):e1010243, 2023.

[29] J Sjovall, D Westerlund, and G Alvan. Renal excretion of intravenously infused amoxycillin and ampicillin. British Journal of Clinical Pharmacology, 19(2):191–201, 1985.

[30] Chad W Stratilo, Scott Jager, Melissa Crichton, and James D Blanchard. Evaluation of liposomal ciprofloxacin formulations in a murine model of anthrax. PloS one, 15(1):e0228162, 2020.

[31] Alexis Tabah, Jan De Waele, Jeffrey Lipman, Jean Ralph Zahar, Menino Osbert Cotta, Greg Barton, Jean-Francois Timsit, and Jason A Roberts. The ADMIN-ICU survey: a survey on antimicrobial dosing and monitoring in ICUs. Journal of Antimicrobial Chemotherapy, 70(9):2671–2677, 2015.

[32] Nassim Nicholas Taleb and Jeffrey West. Working with convex responses: Antifragility from finance to oncology. Entropy, 25(2):343, 2023.

[33] Lindsey HM Te Brake, Veronique de Jager, Kim Narunsky, Naadira Vanker, Elin M Svensson, Patrick PJ Phillips, Stephen H Gillespie, Norbert Heinrich, Michael Hoelscher, Rodney Dawson, et al. Increased bactericidal activity but dose-limiting intolerability at 50 mg· kg-1 rifampicin. European Respiratory Journal, 58(1), 2021.

[34] Pirommas Techitnutsarut and Farida Chamchod. Modeling bacterial resistance to antibiotics: bacterial conjugation and drug effects. Advances in Difference Equations, 2021(1):290, 2021.

[35] U.S. Food and Drug Administration. Ampicillin for Injection, USP. https://www.fda.gov/media/127633/download, 2023. FDA Drug Label.

[36] Bruno Van Herendael, Axel Jeurissen, Paul M Tulkens, Erika Vlieghe, Walter Verbrugghe, Philippe G Jorens, and Margareta Ieven. Continuous infusion of antibiotics in the critically ill: The new holy grail for beta-lactams and vancomycin? Annals of Intensive Care, 2:1–12, 2012.

[37] Jakko Van Ingen, Rob E Aarnoutse, Peter R Donald, Andreas H Diacon, Rodney Dawson, Georgette Plemper van Balen, Stephen H Gillespie, and Martin J Boeree. Why do we use 600 mg of rifampicin in tuberculosis treatment? Clinical Infectious Diseases, 52(9):e194–e199, 2011.

[38] Paul G Williams, Alexis Tabah, Menino Osbert Cotta, Indy Sandaradura, Salmaan Kanji, Marc H Scheetz, Sahand Imani, Muhammed Elhadi, Sonia Luque-Pardos, Natalie Schellack, et al. International survey of antibiotic dosing and monitoring in adult intensive care units. Critical Care, 27(1):241, 2023.

[39] Christopher Witzany, Jens Rolff, Roland R Regoes, and Claudia Igler. The pharmacokinetic–pharmacodynamic modelling framework as a tool to predict drug resistance evolution. Microbiology, 169(7):001368, 2023.

