## Supplemental Material for "A General Analytic Approach to Predicting the Best Antibiotic Dosing Regimen"

**F Code for Figures and Numerical Demonstrations** **10**

### A Definition of Parameters

Table A1 contains all the parameters we use and reference in the paper in the order they appear.

| Parameter | Definition |
|---|---|
| $T$ | Single dose treatment length (h) |
| $T_p$ | Pulse length (for step model) (h) |
| AUC | Area under the curve of the antibiotic concentration (mg · h/L) |
| $D_1$ | Constant-concentration dose level (mg/L) |
| $D_2$ | Pulse (step) dose level (mg/L) |
| $n$ | Hill coefficient (steepness) of Hill function (unitless). Regoes et. al uses $\kappa$ [5], but we use $n$ throughout |
| $G_{\max}$ | Maximum net growth rate of bacteria ($h^{-1}$), also called $\psi_{\max}$ in Regoes et al. [5] |
| $t_{\text{half}}$ | Half-life of the antibiotic (h) |
| $\lambda$ | Exponential decay constant ($\lambda = \ln(2)/t_{\text{half}}$) ($h^{-1}$) |
| $A_0$ | Initial drug concentration (mg/L) |
| $\psi_{\min}$ | Minimum net growth rate of bacteria ($h^{-1}$) |
| MIC | Minimum inhibitory concentration of the antibiotic (mg/L) |
| $V$ | Volume of distribution of the antibiotic (L) |

Table A1: List of parameters referenced throughout, in order of appearance.

### B Proofs of Concavity Results

#### B.1 Proofs of Preliminary Lemmas

**Lemma 1.** *Consider a function $g : [0, \infty) \to \mathbb{R}$.*

1. *If $g$ is strictly concave down and $g(0) = 0$, then $\frac{1}{x} g(x)$ is strictly decreasing.*
2. *If $g$ is strictly concave up and $g(0) = 0$, then $\frac{1}{x} g(x)$ is strictly increasing.*

*Proof.*

1. By the definition of strictly concave down, $\forall w \neq z \in [0, \infty)$ and $\alpha \in (0, 1)$, we have

$$(1 - \alpha)g(z) + \alpha g(w) < g((1 - \alpha)z + \alpha w).$$

Fix $x < y \in (0, \infty)$. Note $\frac{x}{y} \in (0,1)$. So the inequality remains true for $z = 0$, $w = y$, and $\alpha = \frac{x}{y}$, which, since $g(0) = 0$, gives us

$$\frac{x}{y}g(y) < g\left(\frac{x}{y}y\right)$$
$$\frac{1}{y}g(y) < \frac{1}{x}g(x)$$

as needed.

2. By the definition of strictly concave up, $\forall w \neq z \in [0, \infty)$ and $\alpha \in (0,1)$, we have

$$(1-\alpha)g(z) + \alpha g(w) > g((1-\alpha)z + \alpha w).$$

Fix $x < y \in (0, \infty)$. Note $\frac{x}{y} \in (0,1)$. So the inequality remains true for $z = 0$, $w = y$, and $\alpha = \frac{x}{y}$, which, since $g(0) = 0$, gives us

$$\frac{x}{y}g(y) > g\left(\frac{x}{y}y\right)$$
$$\frac{1}{y}g(y) > \frac{1}{x}g(x)$$

as needed. □

**Lemma 2.** *Given three non-colinear points* $(x_1, y_1), (x_2, y_2), (x_3, y_3) \in \mathbb{R}^2$, *exactly one of the following is true:*

*i. There exists a concave up function* $f$ *satisfying* $f$ *continuous and* $f(x_i) = y_i$ *for* $i = 1, 2, 3$.

*ii. There exists a concave down function* $f$ *satisfying* $f$ *continuous and* $f(x_i) = y_i$ *for* $i = 1, 2, 3$.

*Proof.* Fix three non-colinear points $(x_1, y_1), (x_2, y_2), (x_3, y_3) \in \mathbb{R}$ and assume $x_1 < x_2 < x_3$. Define

$$m_1 = \frac{y_2 - y_1}{x_2 - x_1}, \quad m_2 = \frac{y_3 - y_2}{x_3 - x_2}.$$

Now define

$$f(x) = \begin{cases} m_1(x - x_2) + y_2, & x \leqslant x_2 \\ m_2(x - x_2) + y_2, & x \geqslant x_2. \end{cases}$$

Note $f(x_i) = y_i$ for $i = 1, 2, 3$. Since the points are not colinear, $m_1 \neq m_2$, so exactly one must be true:

1. If $m_1 > m_2$, then $f$ is concave down.

2. If $m_1 < m_2$, then $f$ is concave up.

Now, WLOG, assume $m_1 > m_2$. We claim that no concave up curve can be drawn through the three points. Consider

$$g(x) = \frac{y_3 - y_1}{x_3 - x_1}(x - x_1) + y_1$$

which is the line that passes through $(x_1, y_1)$ and $(x_3, y_3)$. Note $x_1 < x_2 < x_3$ and $g(x_2) < y_2$, so any curve which passes through all three points will contradict the definition of concave up. □

#### B.2 Second Derivative of Hill Function

We begin with the assumptions set up in Section 2.2.3:

$$H(y) = c\frac{y^n}{1 + y^n}, \qquad y \in [0, \infty).$$

Then by the quotient rule, we have

$$H'(y) = c\frac{(1 + y^n)(ny^{n-1}) - (y^n)(ny^{n-1})}{(1 + y^n)^2} = c\frac{ny^{n-1}}{(1 + y^n)^2}.$$

By a second application of the quotient rule, we have

$$H''(y) = c\frac{(1 + y^n)^2 \left(n(n-1)y^{n-2}\right) - (ny^{n-1})(2(1 + y^n)(ny^{n-1}))}{(1 + y^n)^4}.$$

Note since $y \geqslant 0$, we have $y^n \geqslant 0$ for $n > 0$ so $1 + y^n > 0$. If $y = 0$ then $H''(0) = 0$, so assume $y > 0$. Then

$$\begin{aligned}
H''(y) = 0 &\iff (1 + y^n)^2 \left(n(n-1)y^{n-2}\right) = 2n^2 y^{2n-2}(1 + y^n) \\
&\iff (1 + y^n)(n(n-1)y^{n-2}) = 2n^2 y^{2n-2} \\
&\iff (n-1)(1 + y^n) = 2ny^n \\
&\iff n - 1 - y^n = ny^n \\
&\iff (n-1)y^{-n} = n + 1
\end{aligned}$$

which is the condition from Section 2.2.3.

### C Proof of Theorem 3

**Theorem 3.** *Given a constant AUC and dose interval $T$, the following hold true when comparing drug regimens in the decay model:*

*i. If the dose response curve is concave up, then the CC regimen performs better than the periodic regimen.*

*ii. If the dose response curve is concave down, then the periodic regimen performs better than the CC regimen.*

Theorem 3 is a special case of Proposition 1, whose statement and proof are presented below. Section C.1 discusses future limitations, context, and future directions using more general pharmacokinetic models to extend the results of this paper.

**Proposition 1.** *Given a constant AUC, treatment length $L > 0$, and any non-negative, bounded, piecewise-continuous, non-constant concentration profile $A_2$ such that $\int_0^L A_2(t)dt = \mathrm{AUC}$, the following hold true when comparing $A_2$ to the CC regimen with the same AUC:*

*i. If the dose response curve is concave up, then the CC regimen performs better than the regimen with concentration profile $A_2$.*

*ii. If the dose response curve is concave down, then the regimen with concentration profile $A_2$ performs better than the CC regimen.*

*Proof.* Fix any non-negative, bounded, piecewise-continuous, non-constant concentration profile $A_2$ on $[0, L]$ such that $\int_0^L A_2(t)dt = \mathrm{AUC}$. Then the CC regimen with the same AUC has drug concentration $A_1(t) = D_C = \mathrm{AUC}/L$ given over the interval $[0, L]$. Under the standard growth rate model, we have

$$\frac{db}{dt} = R\left(A(t)\right) \cdot b(t)$$

where $R$ is the dose response curve, $b$ is the bacteria population, and we define $B(t) = \ln\left(b(t)\right)$ as before. Then on $[0, L]$ we have

$$B_1'(t) = R\left(A_1(t)\right) = R\left(D_C\right), \quad B_2'(t) = R\left(A_2(t)\right).$$

We will also define $A_{\min} = \min_{[0,L]} A_2(t)$ and $A_{\max} = \max_{[0,L]} A_2(t)$ (note $D_C \in (A_{\min}, A_{\max})$) and assume $R$ is concave up on $(A_{\min}, A_{\max})$. We will only prove the result for the concave up case; the result for $R$ concave down on $(A_{\min}, A_{\max})$ follows by flipping all of the inequalities. Since $R$ is concave up on $(A_{\min}, A_{\max})$, we write $R(x) = R_0 - \tilde{R}(x)$ where $\tilde{R}$ is concave down on $(A_{\min}, A_{\max})$ and $\tilde{R}(0) = 0$. We will assume WLOG $B_1(0) = B_2(0) > 0$. We now show $B_2(L) - B_1(L) \geqslant 0$:

$$
\begin{aligned}
B_2(L) - B_1(L) &= \int_0^L B_2'(t)dt - \int_0^L B_1'(t)dt \\
&= \int_0^L (R_0 - \tilde{R}\left(A_2(t)\right))dt - \int_0^L (R_0 - \tilde{R}\left(A_1(t)\right))dt \\
&= \int_0^L \tilde{R}(D_C)dt - \int_0^L \tilde{R}\left(A_2(t)\right)dt \\
&= L\tilde{R}(D_C) - \int_0^L \tilde{R}\left(A_2(t)\right)dt
\end{aligned}
$$

since $\tilde{R}(D_C)$ is constant. Note that $D_C$ is the mean value of $A_2(t)$ over $[0, L]$, i.e.

$$
D_C = \frac{\text{AUC}}{L} = \frac{1}{L}\int_0^L A_2(t)dt.
$$

Since $\tilde{R}$ is concave down on $(A_{\min}, A_{\max})$, by Jensen's inequality, we have

$$
\frac{1}{L}\int_0^L \tilde{R}\left(A_2(t)\right)dt \leqslant \tilde{R}\left(\frac{1}{L}\int_0^L A_2(t)dt\right) = \tilde{R}(D_C).
$$

If we substitute this into our expression for $B_2(L) - B_1(L)$, we get

$$
B_2(L) - B_1(L) \geqslant L\tilde{R}(D_C) - L\tilde{R}(D_C) = 0.
$$

Moreover, if $\tilde{R}$ is strictly concave down on $(A_{\min}, A_{\max})$, then the inequality in Jensen's inequality is strict so $B_2(L) - B_1(L) > 0$.

□

#### C.1 Discussion of More General Pharmacokinetic Models

One limitation in the main text of this work is that we consider two very simple pharmacokinetic models for periodic dosing regimens, the "Step PK" model from Section 2.2.2 which does not take into account any ADME processes (absorption, distribution, metabolism, excretion) and the "Decay PK" model from Section 2.3 which incorporates first-order elimination of the drug. However, real pharmacokinetic profiles can be much more complicated, often requiring multi-compartment models [1] to accurately capture the drug concentration in the target tissue over time to account for effects such as saturation of PK processes [3]. While Proposition 1 provides a general result for any bounded, measurable concentration profile, in this work we always added a second step between Theorems 2 and 3 and their applications to real dose response curves which may exhibit more than one concavity. For these cases, the additional step translating mono-convexity results to "mixed concavity" models incorporates the specific features of the pharmacokinetic model being used.

For example, with the step model, sometimes the concavity of dose response curve is not the same as the concavity of the function we draw through the discrete points $(D_C, R(D_C))$, $(D_P, R(D_P))$, and $(0,0)$. This was not an issue in the decay model because the codomain of $R\left(A_0 e^{-\lambda t}\right)$ is continuous, meaning the concavity of $\{(x, R(x)) : x \in [D_{\min}, D_{\max}]\}$ (note $D_C \in (D_{\min}, D_{\max})$) is the same as the concavity of $R$ on that interval.

So while Proposition 1 holds for any pharmacokinetic model, the particular features of the PK model impact how the results of Proposition 1 can be applied to real dose response curves. Future work could extend the results of this paper to more complicated pharmacokinetic models, such as those incorporating multi-compartment dynamics, saturable processes, or auto- (self-)inhibition of a drug's own metabolism [1, 3].

### D Explicit Extension of Hill Functions to Mono-convex Functions

In this section, we show explicitly that the results of Theorem 3 can be applied to Hill functions with mixed concavity. First, recall three things: our dose response curve $R$ is given by $R(x) = G_{\max} - H(x)$ where $H(x) = c\frac{x^n}{k^n + x^n}$; when the Hill coefficient $n \in (0,1]$, $H$ is concave down, and when $n > 1$, we have an inflection point at $\tilde{D} = k\left(\frac{n-1}{n+1}\right)^{\frac{1}{n}}$ and $H$ is concave down to the left of $\tilde{D}$ and concave up to the right of $\tilde{D}$; when considering a periodic decay regimen with its associated CC regimen of the same AUC, the constant drug concentration $D_C = \frac{A_0}{\lambda T}(1 - e^{-\lambda T})$, and we will notate our minimum and maximum drug concentrations of the periodic dose as

$$D_{\min} = A_0 e^{-\lambda T}, \quad D_{\max} = A_0.$$

Now note

$$D_{\min} < D_C < D_{\max}.$$

This means we need to consider the concavity of the dose response curve on the *interval* of doses $\mathcal{D} := (D_{\min}, D_{\max})$ to determine which regimen performs better. Since we are considering a contiguous interval instead of a discrete set of doses (unlike the step model where $\mathcal{D}$ was a set of three dose sizes), we will never be in a case where the dose response curve is, for example, concave up while simultaneously being able to draw a concave down curve through all the points in $\mathcal{D}$ since $\mathcal{D}$ is connected for the decay model. We have the following possible cases:

I. If $n \in (0,1]$, then $H$ is concave down, and since $H(0) = 0$, by direct application of Theorem 3, the CC regimen performs better than the periodic regimen.

II. If $n > 1$ and $D_{\max} \leqslant \tilde{D}$, then $H$ is concave up on $(0, D_{\max})$, so we can extend $H$ to a concave up function on all of $(0, \infty)$ by defining

$$\tilde{H}(x) = H(x) \cdot \chi_{[0, D_{\max}]} + \Omega(x) \cdot \chi_{[D_{\max}, \infty)} \tag{D1}$$

where

$$\Omega(x) = H'(D_{\max})(x - D_{\max}) + H(D_{\max}), \quad x \in [D_{\max}, \infty) \tag{D2}$$

and since $\tilde{H}(0) = 0$, the periodic regimen performs better than the CC regimen.

III. If $n > 1$ and $D_{\min} > \tilde{D}$, then $H$ is concave down on $(D_{\min}, D_{\max})$. Define

$$\tilde{H}(x) := \Omega(x) \cdot \chi_{[0,D_{\min}]} + H(x) \cdot \chi_{[D_{\min},\infty]}$$

where

$$\Omega(x) := \frac{H(D_{\min})}{D_{\min}} x, \quad x \in [0, D_{\min}].$$

Then we can check the concavity of $\tilde{H}$ by calculating $H'(D_{\min})$ and comparing it to the value of $\frac{H(D_{\min})}{D_{\min}}$. We will have two cases:

(a) If $H'(D_{\min}) \leqslant \frac{H(D_{\min})}{D_{\min}}$, then $\tilde{H}$ is concave down and $\tilde{H}(0) = 0$, so by Theorem 3, the CC regimen performs better than the periodic regimen.

(b) If $H'(D_{\min}) > \frac{H(D_{\min})}{D_{\min}}$, then we must translate $H$ up by $K$; call the translation $H_K(x) = H(x) + K$. We choose $K$ such that $H'_K(D_{\min}) \leqslant \frac{H_K(D_{\min})}{D_{\min}}$. We then define

$$\tilde{H}_K(x) := \Omega(x) \cdot \chi_{[0,D_{\min}]} + H_K(x) \cdot \chi_{[D_{\min},\infty]}$$

where

$$\Omega(x) := \frac{H_K(D_{\min})}{D_{\min}} x, \quad x \in [0, D_{\min}].$$

Then $\tilde{H}_K$ is concave down and $\tilde{H}_K(0) = 0$, so by Theorem 3, the CC regimen performs better than the periodic regimen. We can do this because vertical translations preserve concavity, and we can rewrite

$$R(x) = (G_{\max} + K) - H_K(x).$$

IV. If $n > 1$ and $D_{\min} < \tilde{D} < D_{\max}$, then with the theory we have built up in this paper, we do not know which regimen will perform better.

Cases (II), (IIIa), and (IIIb) are illustrated in Figure D1.

### E Completed Details for Numerical Simulations

#### E.1 Dose Response Curves of Rifampin, Ciprofloxacin, Streptomycin, and Tetracycline

Figure E2 shows the dose response curves of rifampin, ciprofloxacin, streptomycin, and tetracycline against *E. coli* in vitro. The dose response curves are parametrized from data in [5] and are on a $\log(x)$ scale. Note this means that for the antbiotics with $n > 1$, the inflection point does not appear at the graph's visual inflection point. A comparison we made previously was that the inflection point of ciprofloxacin is much lower than the inflection point of rifampin, which is clear from this figure.

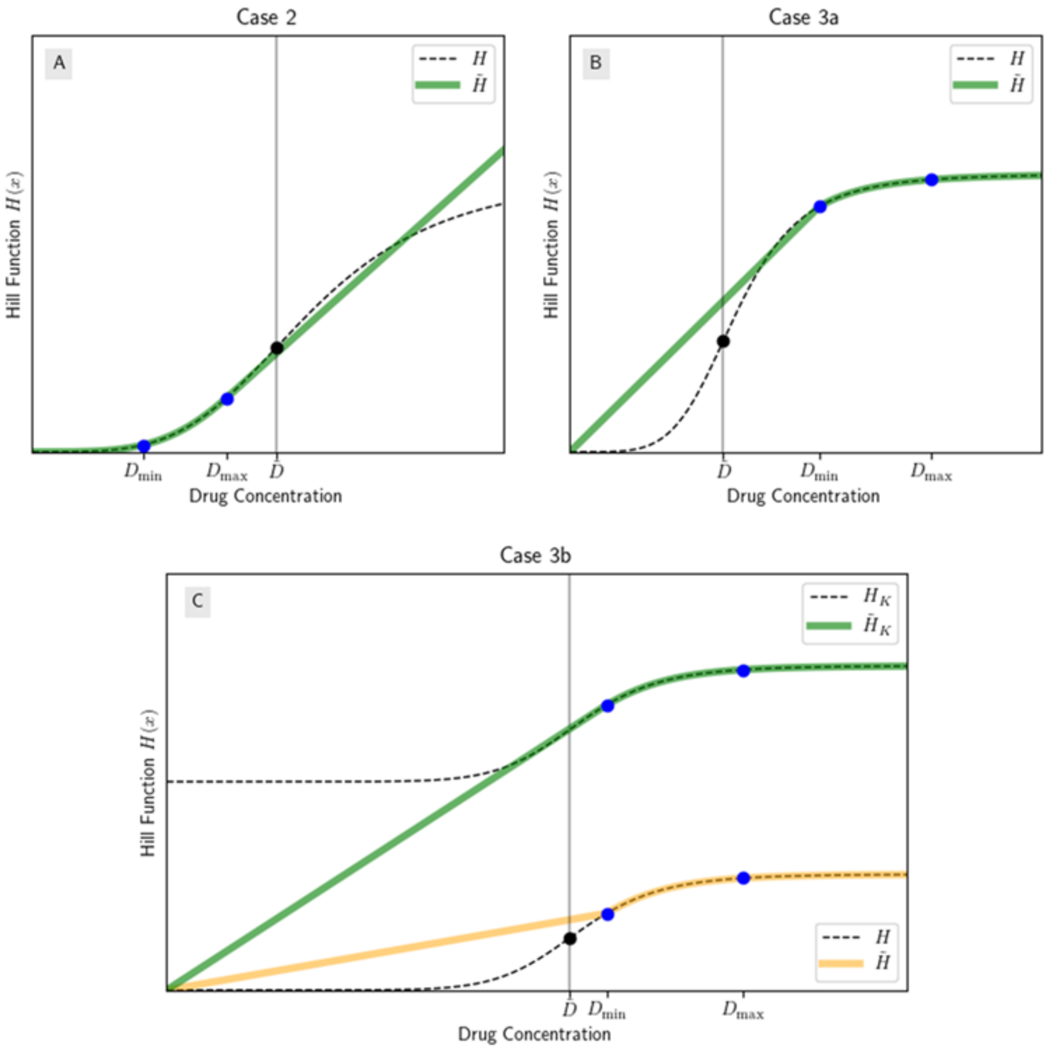

Figure D1: Cases (II), (IIIa), and (IIIb) from Section 2.3. Each case has Hill Coefficient $n > 1$ so each is an example of a Hill function with mixed concavity. (A) $D_{\max} < \tilde{D}$, so we extend $H$ to a concave up function on all of $(0, \infty)$ by defining a linear function on $(D_{\max}, \infty)$; (B) $D_{\min} > \tilde{D}$ and $H'(D_{\min}) \leqslant \frac{H(D_{\min})}{D_{\min}}$, so we extend $H$ to a concave down function on all of $(0, \infty)$ by defining a linear function on $(0, D_{\min})$; (C) $D_{\min} > \tilde{D}$ and $H'(D_{\min}) > \frac{H(D_{\min})}{D_{\min}}$, so we extend $H$ to a concave down function by translating it up, then defining a linear function on $(0, D_{\min})$.

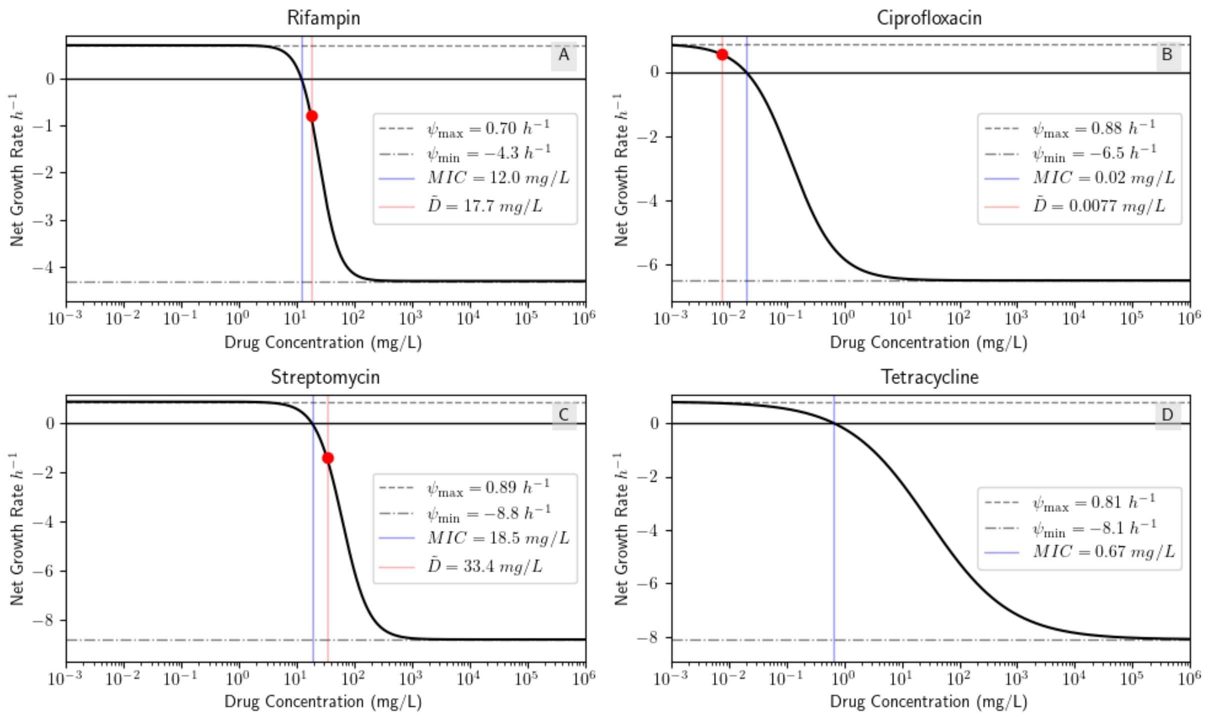

Figure E2: Dose response curves of rifampin, ciprofloxacin, streptomycin, and tetracycline. Data from [5].

#### E.2 Full Regimen Details for Ciprofloxacin and Rifampin Simulations

Table E2 and Table E3 contain the full details of the dosing regimens we defined in Section 3.2 as well as each drug's PD parameters. Figures E3 and E4 contain the bacteria population curves for each ciprofloxacin and rifampin regimen.

### F Code for Figures and Numerical Demonstrations

The code used to generate Figure 1 as well as the numerical results in this paper is available in the following Github repository:
https://github.com/leahchilders/GeneralAntibioticDosing. The code is written in a Jupyter Notebook using Python 3.12.8 and uses the SciPy library for numerical integration. The code is designed to be user-friendly and can be used to analyze any antibiotic data with a selection of PD models. The user can choose a PD model and input their own PK and PD parameters. The code outputs, among other things, the value of $\tilde{D}$, an analytic and numerical analysis of the concavity of the Hill function as performed in this paper, and plots of the dose response curve, the treatment regimens, and the bacteria population curves. The code to reproduce Figure 1 is also provided in a separate Jupyter

| **Parameter** | **Regimen 1** | **Regimen 2** | **Regimen 3** | **Regimen 4** |
|---|---|---|---|---|
| Half-life (h) | 4 | | | |
| Volume of distribution (L) | 210 | | | |
| Periodic dose size (mg) | 100 | 1 | 30 | 2.5 |
| Periodic max conc. (mg/L) | 0.48 | 0.0048 | 0.14 | 0.012 |
| Periodic min conc. (mg/L) | 0.17 | 0.0006 | 0.0022 | 0.0015 |
| Dose period (h) | 6 | 12 | 24 | 12 |
| AUC (mg · h/L) | 1.78 | 0.024 | 0.81 | 0.06 |
| Constant conc. (mg/L) | 0.3 | 0.002 | 0.034 | 0.005 |
| $\psi_{\text{max}}$ ($h^{-1}$) | 0.88 | | | |
| $\psi_{\text{min}}$ ($h^{-1}$) | $-6.5$ | | | |
| MIC (mg/L) | 0.02 | | | |
| $n$ | 1.1 | | | |
| $\tilde{D}$ (mg/L) | 0.0077 | | | |
| Periodic bac. end value | $4.46 \times 10^{-10}$ | $1.48 \times 10^{6}$ | $1.97 \times 10^{-2}$ | $3.00 \times 10^{5}$ |
| Constant bac. end value | $2.34 \times 10^{-10}$ | $1.50 \times 10^{6}$ | $1.67 \times 10^{-4}$ | $3.04 \times 10^{5}$ |
| Which regimen performed better? | Continuous | Periodic | Continuous | Periodic |

Table E2: Four ciprofloxacin regimens and the bacteria population value at the end of each treatment interval. Constant concentration is calculated as the AUC of the periodic dose divided by the dose period. Half-life from [4], volume of distribution from [2], PD parameters from [5]. End bacteria population values are given as a percentage of the initial bacteria population

Notebook.

| **Parameter** | **Reg. 1** | **Reg. 2** | **Reg. 3** | **Reg. 4** | **Reg. 5** |
|---|---|---|---|---|---|
| Half-life (h) | 2.5 | | | | |
| Volume of dist. (L) | 53.2 | | | | |
| Per. dose size (mg) | 7000 | 3100 | 3000 | 1400 | 500 |
| Per. max conc. (mg/L) | 131.58 | 58.27 | 56.39 | 26.32 | 9.40 |
| Per. min conc. (mg/L) | 24.93 | 2.09 | 2.02 | 0.94 | 0.34 |
| Dose period (h) | 6 | 12 | 12 | 12 | 12 |
| AUC ($mg \cdot h/L$) | 384.66 | 202.62 | 196.09 | 91.51 | 32.68 |
| Cont. conc. (mg/L) | 64.11 | 16.89 | 16.34 | 7.63 | 2.72 |
| $\psi_{max}$ ($h^{-1}$) | 0.7 | | | | |
| $\psi_{min}$ ($h^{-1}$) | $-4.3$ | | | | |
| MIC (mg/L) | 12.0 | | | | |
| $n$ | 2.5 | | | | |
| $\tilde{D}$ (mg/L) | 17.67 | | | | |
| Per. end bac. value | $7.93 \times 10^{-8}$ | $4.12 \times 10^{-2}$ | $6.98 \times 10^{-2}$ | $1.73 \times 10^{3}$ | $2.41 \times 10^{5}$ |
| Cont. end bac. value | $8.04 \times 10^{-9}$ | $2.76 \times 10^{-2}$ | $7.25 \times 10^{-2}$ | $2.24 \times 10^{4}$ | $3.50 \times 10^{5}$ |
| Which reg. performed better? | Continuous | Continuous | Periodic | Periodic | Periodic |

Table E3: Five rifampin regimens and the bacteria population value at the end of each treatment interval. Constant concentration is calculated as the AUC of the periodic dose divided by the dose period. Half-life from [6], volume of distribution from [7], Regoes parameters from [5]. End bacteria values are given as a percentage of the initial bacteria population.

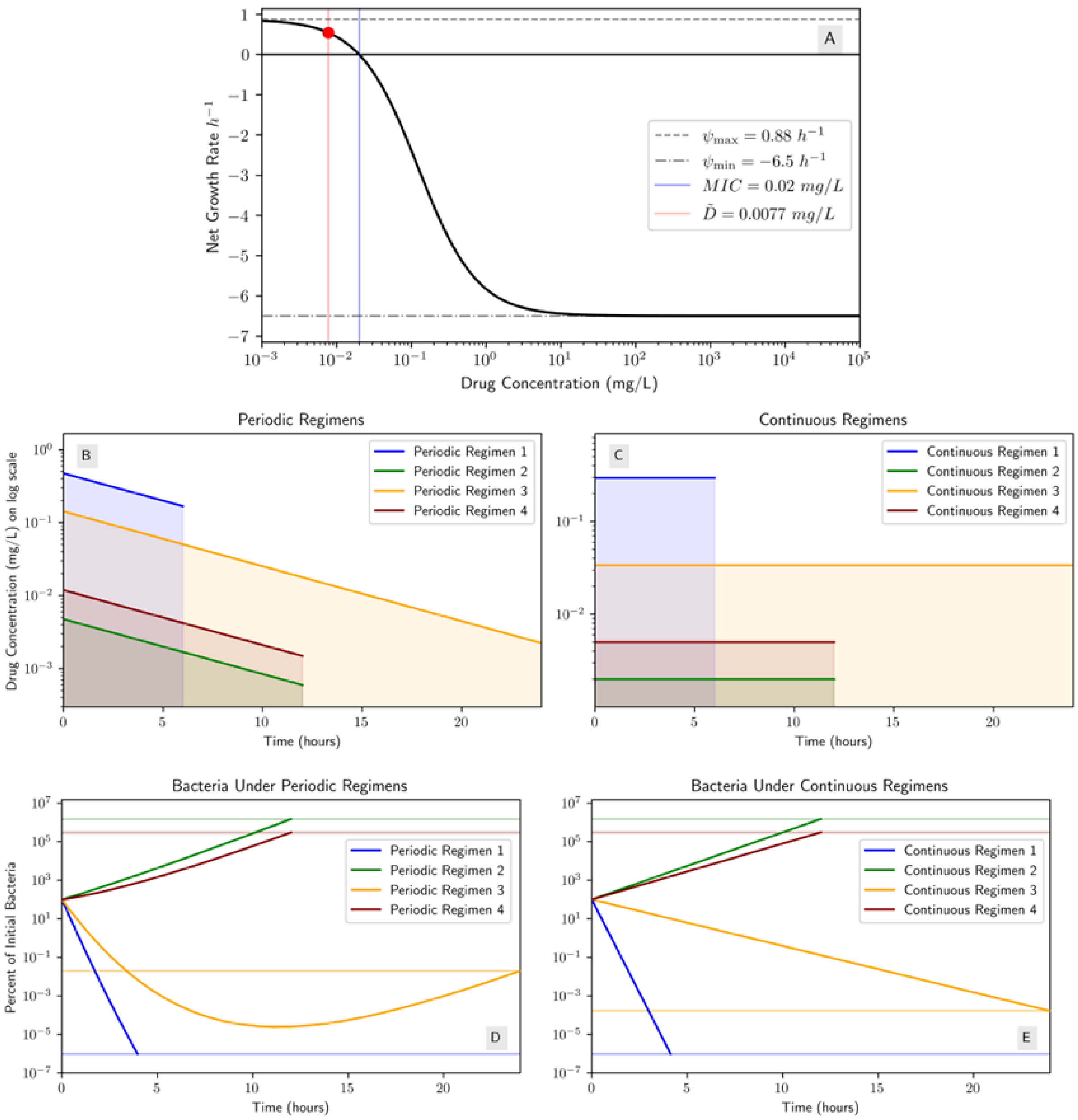

Figure E3: Bacteria population curves for ciprofloxacin regimens from Table E2. (A) Dose response curve for ciprofloxacin [5]; (B) periodic dosing regimens described in Table E2; (C) the periodic regimens' respective CC regimens (i.e. CC regimen $i$ has the same AUC as periodic regimen $i$); (D) the bacteria population under each periodic regimen; (E) the bacteria population under each CC regimen.

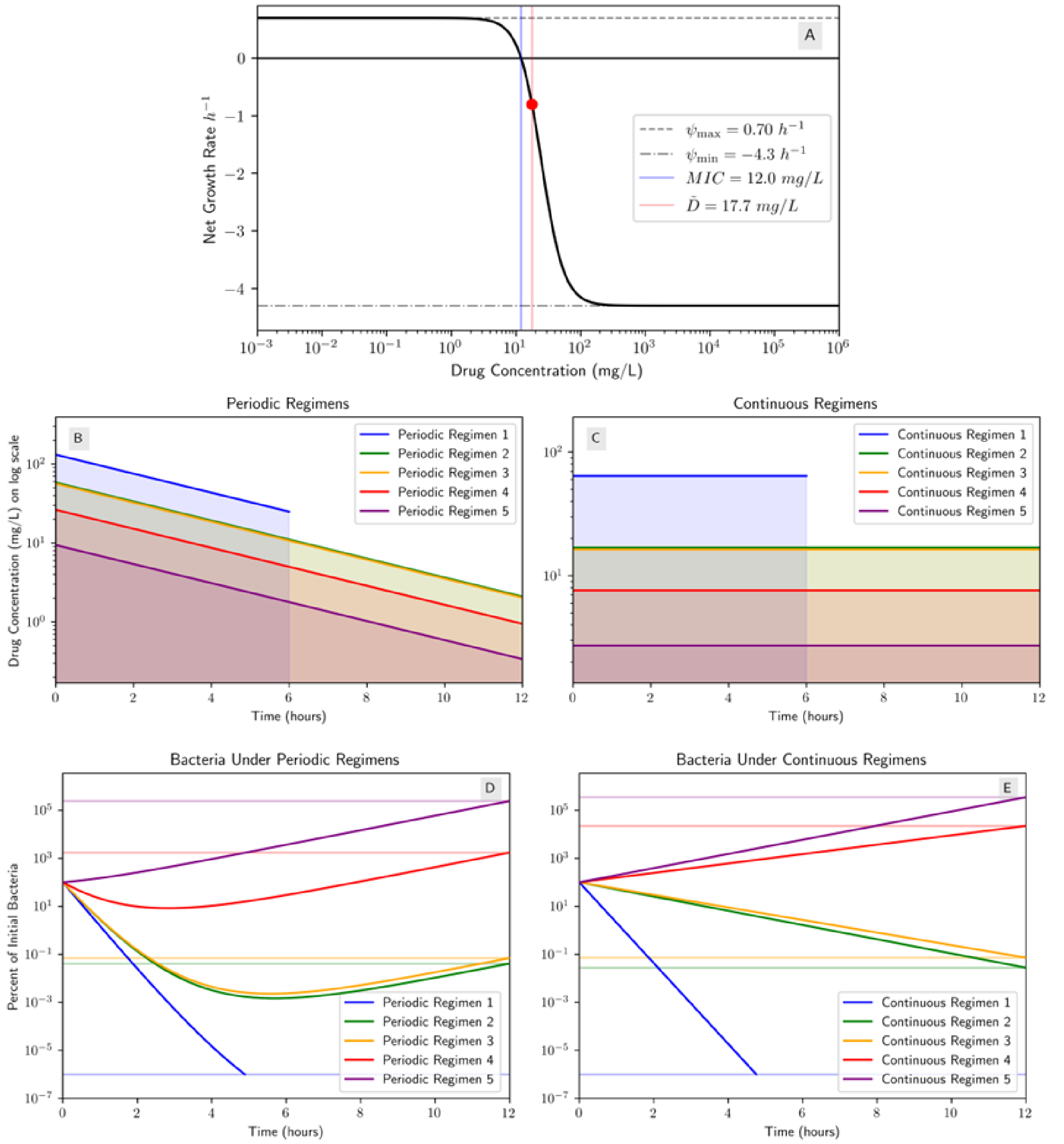

Figure E4: Bacteria population curves for rifampin regimens from Table E3. (A) Dose response curve for rifampin [5]; (B) periodic dosing regimens described in Table E3; (C) the periodic regimens' respective CC regimens (i.e. CC regimen $i$ has the same AUC as periodic regimen $i$); (D) the bacteria population under each periodic regimen; (E) the bacteria population under each CC regimen.

### References

[1] Khaled Abduljalil, Martina Kinzig, Juürgen Bulitta, Stefan Horkovics-Kovats, Fritz Sörgel, Michael Rodamer, and Uwe Fuhr. Modeling the autoinhibition of clarithromycin metabolism during repeated oral administration. *Antimicrobial Agents and Chemotherapy*, 53(7):2892–2901, 2009.

[2] Evan J Begg, Richard A Robson, Darren A Saunders, Garry G Graham, Rona C Buttimore, Alister M Neill, and G Ian Town. The pharmacokinetics of oral fleroxacin and ciprofloxacin in plasma and sputum during acute and chronic dosing. *British Journal of Clinical Pharmacology*, 49(1):32–38, 2000.

[3] Femke de Velde, Brenda CM de Winter, Birgit CP Koch, Teun van Gelder, and Johan W Mouton. Non-linear absorption pharmacokinetics of amoxicillin: consequences for dosing regimens and clinical breakpoints. *Journal of Antimicrobial Chemotherapy*, 71(10):2909–2917, 2016.

[4] Bayer HealthCare Pharmaceuticals Inc. CIPRO- ciprofloxacin hydrochloride tablet, film coated. `https://dailymed.nlm.nih.gov/dailymed/fda/fdaDrugXsl.cfm?setid=888dc7f9-ad9c-4c00-8d50-8ddfd9bd27c0&type=display`, 01 2023.

[5] Roland Regoes, Camilla Wiuff, Renata Zappala, Kim Garner, Fernando Baquero, and Bruce Levin. Pharmacodynamic Functions: a Multiparameter Approach to the Design of Antibiotic Treatment Regimens. *Antimicrobial Agents and Chemotherapy*, 48:3670–6, 10 2004.

[6] Jakko Van Ingen, Rob E Aarnoutse, Peter R Donald, Andreas H Diacon, Rodney Dawson, Georgette Plemper van Balen, Stephen H Gillespie, and Martin J Boeree. Why do we use 600 mg of rifampicin in tuberculosis treatment? *Clinical Infectious Diseases*, 52(9):e194–e199, 2011.

[7] Justin J Wilkins, Radojka M Savic, Mats O Karlsson, Grant Langdon, Helen McIlleron, Goonaseelan Pillai, Peter J Smith, and Ulrika SH Simonsson. Population pharmacokinetics of rifampin in pulmonary tuberculosis patients, including a semimechanistic model to describe variable absorption. *Antimicrobial Agents and Chemotherapy*, 52(6):2138–2148, 2008.
